# Modelling, characterization of data-dependent and process-dependent errors in DNA data storage

**DOI:** 10.1101/2021.07.17.452779

**Authors:** Yixin Wang, Md Noor-A-Rahim, Erry Gunawan, Yong Liang Guan, Chueh Loo Poh

## Abstract

**Motivation:** Using DNA as the medium to store information has recently been recognized as a promising solution for long-term data storage. While several system prototypes have been demonstrated, the error characteristics in DNA data storage are discussed with limited content. Due to the data and process variations from experiment to experiment, the error variation and its effect on data recovery remain to be uncovered. To close the gap, we systematically investigate the storage channel, i.e., error characteristics in the storage process.

**Results:** We first propose a new concept named sequence corruption to unify the error characteristics into the sequence level, easing the channel analysis. Then we derived the formulations of the data imperfection at the decoder including both sequence loss and sequence corruption, revealing the decoding demand and monitoring the data recovery. Furthermore, we extensively explored several data-dependent unevenness observed in the base error patterns and studied a few potential factors and their impacts on the data imperfection at the decoder both theoretically and experimentally. The results presented here introduce a more comprehensive channel model and offer a new angle towards the data recovery issue in DNA data storage by further elucidating the error characteristics of the storage process.

## 1 Introduction

The explosion of data has driven scientists to explore new technologies to store information. In recent years, owing to the superior properties like extremely high physical density and preservation duration, using DNA molecules as the data storage medium has drawn a rising attention (Church *et al.*, 2012; Goldman *et al.*, 2013; Grass *et al.*, 2015; Yazdi *et al.*, 2015; Bornholt *et al.*, 2016; Blawat *et al.*, 2016; Yazdi *et al.*, 2017; Erlich and Zielinski, 2017; Organick *et al.*, 2018; Choi *et al.*, 2019; Wang *et al.*, 2019b). In a typical DNA data storage system, the basic data unit is a DNA strand that represents a string of nucleotide bases consisting of Adenine (A), Thymine (T), Cytosine (C), and Guanine (G). Data writing in DNA data storage is performed by encoding the digital information into an assemble of DNA sequences. Taking the encoded DNA sequences as the reference, corresponding DNA molecules are synthesized and very often the number of molecule copies (e.g., copies of oligo) of each reference sequence varies. The synthesized DNA can then be stored and sequenced during which several random processes are involved, leading to *sequence loss* at the decoder (Erlich and Zielinski, 2017; Chen *et al.*, 2020). Here, sequence loss refers to the loss of all copies/reads of the reference sequence. In other words, if all sequence data of one reference sequence could not be found at the decoder, this reference sequence is considered as lost.

Besides the sequence loss, *base error* is the other type of error in DNA data storage. While the general base error statistics have been reported (Organick *et al.*, 2018; Heckel *et al.*, 2019) and the sequence loss has been studied (Chen *et al.*, 2020; Heckel *et al.*, 2019), the overall impact of these two types of errors, i.e., base level and sequence level, on the decoder, remains unclear. To unify the error characteristics and ease the analysis at the decoder, we introduce the concept of *sequence corruption*, transmitting the base type error to sequence type error by incorporating the effects of physical redundancy (i.e., multiple sequence copies at the receiver/sequencer) and post-processing method into the channel model. Different from sequence loss, sequence corruption refers to the failure of reconstructing the reference sequence from its erroneous copies (i.e., with base errors) at the receiver, which is the direct consequence of the base error. Theoretically, the sequence corruption rate collectively depends on the base error rate in the received copies of the reference sequence, the received copy counts of the reference sequence, and the post-processing methods of reconstructing the reference sequence. From another perspective, this new concept leverages the multi-count physical redundancy feature of DNA data storage, enabling the anticipation of the required logical redundancy in code design with the presence of specific physical redundancy in experiment. As a result, we define the data imperfection at the decoder of DNA data storage channel consisting of sequence loss and sequence corruption. Investigating the characteristic of base errors where some variance might exist is also important since it essentially relates to sequence corruption and can guide the sequence (codeword) design. Meanwhile, many factors, including data structure design, experiment design and computational processing, could affect the degree of data imperfection at the decoder, leading to process-dependent errors. Understanding how the original data (reference sequences) and these factors affect the overall error rate at the decoder could provide insights into several aspects of designing advanced DNA data storage systems, including codec designs, sequence structure designs, experiment designs, and data processing methods.

In this paper, we theoretically formulated the sequence corruption which is cooperatively dictated by the base error statistics, copy counts of reference sequence, and down-stream processing methods. Combining the sequence corruption with the sequence loss, we then quantified the data imperfection by deriving the overall sequence error rate of the DNA data storage channel. The derivation explicitly takes the unevenness in both the count distribution and the error patterns into consideration, revealing distinct data recovery demands in DNA data storage using different sequencing techniques. Furthermore, we investigated the base error properties by analyzing the data from our previous work (Wang *et al.*, 2019b,c). Specifically, we first looked into the single base error and then analyzed the 2/3/4-mer patterns with different types of errors, i.e., substitutions, insertions, and deletions. We observed that there are profound biases in transitions errors among DNA bases; and certain k-mer patterns (not only homopolymers) are prone to certain type of errors. Lastly, with data collected from two independent experiments and theoretical analysis, we broadly studied the factors that might affect the data integrity in aspects spanning from structure design to biological and analytical handling methods. By conducting the most comprehensive study on the imperfect and uneven data in DNA data storage so far, the results in this work could offer insights and instructions to the design and processing pipeline of more effective and efficient DNA data storage systems.

## 2 Approach and Method

### 2.1 Data flow and errors in DNA data storage

Data are represented in different forms at different stages in the DNA data storage, such as binary stream, DNA sequences, and physical DNA molecules (see Fig. 1A). Binary data are encoded and converted into DNA sequences before sending them to DNA synthesis. At the synthesis stage, the count of the oligos may vary, and the count distribution can be approximated by gamma or normal distribution based on the different synthesis techniques (Chen *et al.*, 2020). Following that, sample might be stored in a distributed fashion to increase data accessibility, where a random process happens. To illustrate, Fig. 1A describes one scenario where physical copies of certain (i.e., purple-colored) reference sequence are all lost, rendering *physical sequence loss* (i.e., 0 physical copy of the reference sequence).

**Fig. 1.**
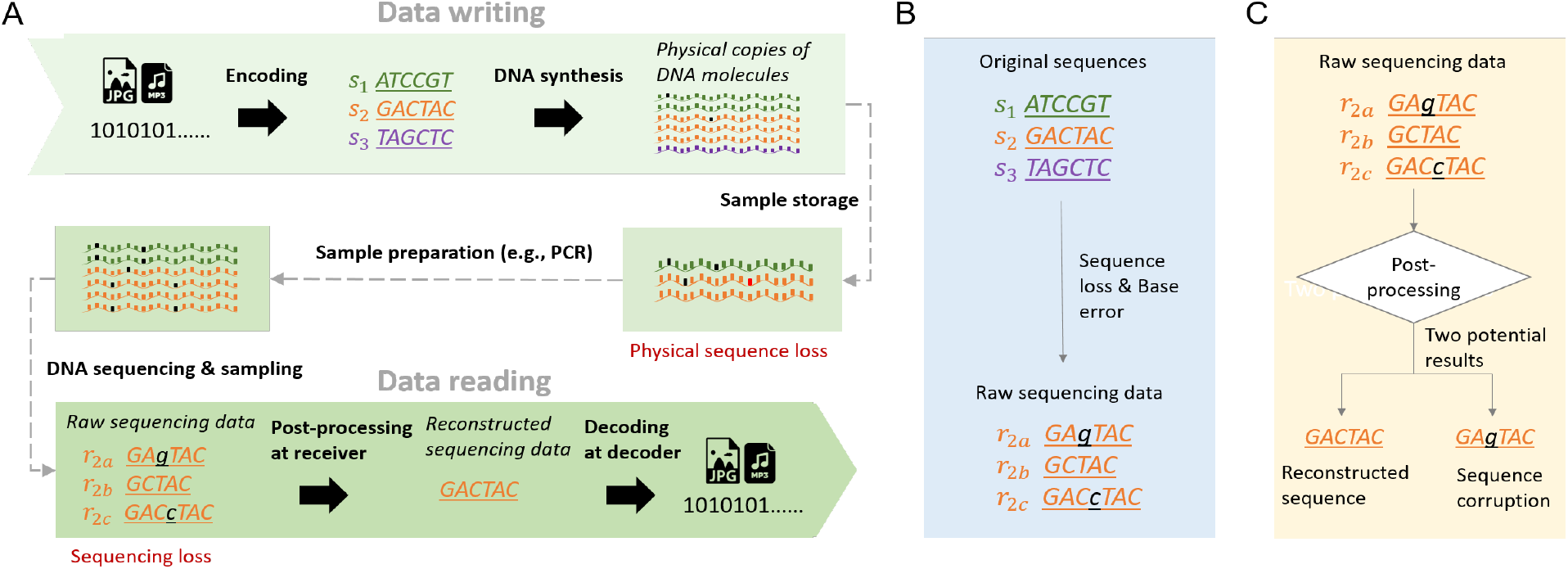
Data flow and error characterization in DNA data storage.

To prepare the stored sample for DNA sequencing, the sample is usually PCR amplified to meet the sequencing requirements. The PCR amplification is another random process, where the count of newly generated molecule follows a binomial distribution with a probability of successful amplification. This process is likely to exacerbate the bias on the count distribution. This biased count distribution usually lead to additional data (sequence) loss at the DNA sequencing stage since the next-generation sequencing process is another round of random sampling (Aird *et al.*, 2011), i.e., the chip only reads certain amounts of molecules from a molecule population. With a population of highly biased distribution, each element may have a different probability of being sampled. Hence, if the sampling size is inadequate (i.e., low sequencing coverage), the Poisson sampling effect would cause another round of data loss. We categorize data loss at this stage as *sequencing loss* (see Fig. 1A).

Apart from sequence loss, base error is the other type of errors in the sequencing data at the receiver. In Fig. 1A and B, hypothesized random base errors are black-colored for illustration. Before sending to the decoder, the received raw sequencing data is usually post-processed for a preliminary data reconstruction as shown in Fig. 1A and C. Note that no standard has been set yet for processing the sequencing data while the processing results given by different processing methods definitely affect how many remained errors that the decoder needs to handle. Fig. 1C shows two potential processing results, i.e., successful reconstructed sequence and sequence corruption, of which the sequence corruption remains to be resolved by the decoder.

### 2.2 Pair-end sequencing and sequence alignment

Next-generation sequencing technologies provide protocols to generate reads from two ends of the DNA strand. These protocols enable the sequencer to recover long DNA sequences given that the sequence length is no longer than twice the read length. Besides, pair-end reading is also recognized to improve the sequencing accuracy due to the overlapping between the pair of reads. In our two previous works, Pair-end 150 (PE150) protocols were used to read DNA oligos with lengths from 190 to 199 (Wang *et al.*, 2019b,c) to make full use of the current synthesis and sequencing technologies. To merge the pair-end short reads into the long reads, several prevalent tools were designed (Zhang *et al.*, 2014; Masella *et al.*, 2012; Magoč and Salzberg, 2011; Liu *et al.*, 2012), among which we used FLASH (Magoč and Salzberg, 2011) to merge PE150 reads (see Supplementary S1). To estimate the base error statistics, the merged reads from PE reads are aligned to their corresponding reference/original sequences using sequence alignment tools. Several tools were devised for sequence alignment (Langmead and Salzberg, 2012; Li *et al.*, 2009; Li and Durbin, 2009), among which we used Bowtie 2 (Langmead and Salzberg, 2012).

## 3 Result

### 3.1 Deriving the overall sequence error rate consisting of sequence loss and sequence corruption

Sequence loss in DNA data storage channel might be due to the physical sequence loss in the sample preparation and storage and/or the sequencing loss in sequencing. We used a model which computationally simulates the whole process of DNA data storage in (Chen *et al.*, 2020) to study the sequence loss at the decoder. By defining *channel coverage* as the average number of reads per reference sequence the decoder receives, we found that when the channel coverage is sufficient, the overall sequence loss rate is lower bounded by the physical sequence loss (see Supplementary S2 Fig. 3). Without loss of generality, the sequence loss is found to be higher when data are sampled from a population with more severe over-dispersion, i.e., smaller coefficient of variation (C.V.) (Supplementary S2 Fig. 4). For insufficient channel coverage scenario (i.e., less than 10x), it was found that the overall sequence loss rate no longer changes linearly with the physical sequence, implying that the sequencing loss dominates the overall dropout rate (Supplementary S2 Fig. 5). We also evaluated the model by fitting it with data from our previous work (Wang *et al.*, 2019c), where the correlation coefficient (i.e., *R*^2^ = 0.96) shows that the sequencing sampling effect is well-simulated (Supplementary S2 Fig. 6). Moreover, by comparing the experimental dilution effect in (Erlich and Zielinski, 2017) with the simulated dilution effect in a modified version of the computational model in (Chen *et al.*, 2020), we found that there is still a notable gap between the experiment and the simulation (see Supplementary S2 and S2 Fig. 7). To further understand the gap, we used three different types of amplification efficiency *p*, i.e., constant, random, and strand-specific random, in the computational model to probe the association between PCR amplification and the source of the gap (see Supplementary S2 Fig. 8). Applying strand-specific randomness in the computational model gives the closest approximation to the experimental results in (Erlich and Zielinski, 2017).

Sequence corruption was underestimated in the existing works (Erlich and Zielinski, 2017; Organick *et al.*, 2018; Chen *et al.*, 2020; Heckel *et al.*, 2019). However. the impact of sequence corruption on the decoding is not trivial when establishing more cost-effective and large-scaled DNA data storage with less accurate synthesis and sequencing technologies where base error rates are higher and the copy counts of the reference sequences are limited at the receiver. In the following, we extensively study the overall sequence error rate by incorporating both sequence loss and sequence corruption.

#### 3.1.1 Simplified derivation of sequence error rate

We start from the simplest formulation in which the copy count is assumed to be even and the base error rate is assumed to be constant, i.e., each base has the same error probability. With these assumptions, there is no sequence loss but only sequence corruption that is stemmed from the base error; and it highly depends on the available copy counts at the receiver which is denoted as channel coverage *η*. Besides, the sequence corruption rates may vary if different post-processing methods are used before decoding. Here, we formulate the sequence error rate for two commonly adapted methods, i.e., non-consensus (or trial-and-error) and consensus (i.e., majority selection at each position). The trial-and-error means one reference sequence is regarded as correctly recovered if at least one read copy of it at the receiver is error-free. The majority selection is a well-known consensus algorithm for generating representative data of clustered data. One reference sequence is regarded as correctly recovered if the representative sequence is error-free. For simplicity, the formulation temporarily assumes binary majority selection at each position.

Illumina and Nanopore sequencing are the two commonly used sequencing techniques in the existing DNA data storage where Nanopore sequencing can sequence longer sequence but provides lower sequencing accuracy. To show how channel coverage affects the sequence error rate in Illumina- and Nanopore-based DNA data storage differently, we applied correspondingly different values to parameters including base error rate *ϵ*, sequence length *M*, to the formulation (see Supplementary S3) and depicted the sequence error rate against the channel coverage. In Fig. 2, all embedded figures in the sub-figures refer to the Nanopore-based while the rest refers to the Illumina-based. It was found that, using Illumina sequencing, the sequence corruption rate decreases drastically with the increase of channel coverage for both non-consensus (Fig. 2A) and consensus cases (Fig. 2B). Specifically, with only ~ 5 channel coverage, the corruption rate could be reduced nearly to 0; and with addition of a consensus processing, the minimum channel coverage is decreased by half, i.e., ~ 2.5. However, for channels using Nanopore sequencing, the corruption rate maintains a high plateau with non-consensus method (embedded figure in Fig. 2A), and decreases much more gradually with the consensus method (embedded figure in Fig. 2B). With consensus algorithm, the corruption rate approaches to 0 for a minimum of ~ 20 coverage. Overall, the observation indicates that in Illumina-based storage, increasing channel coverage (read copy redundancy) could effectively reduce the error rate even without any consensus algorithm while in Nanopore-based storage, only with appropriate consensus algorithm and sufficient coverage, the error rate can be reduced to an acceptable level.

**Fig. 2.**
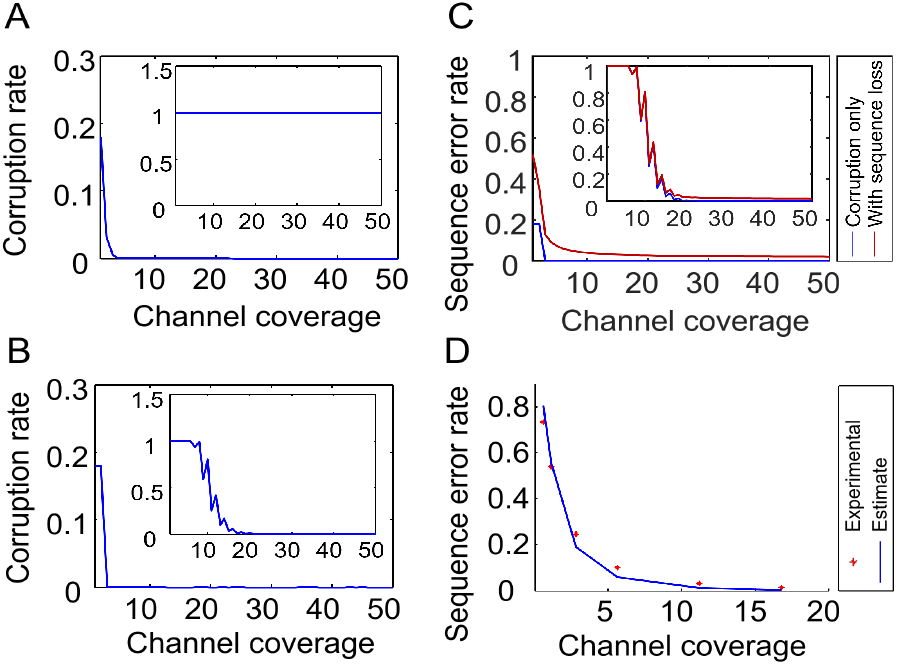
The association between the channel coverage and the sequence corruption and sequence error rate. The major graph in each sub-figure refers to the Illumina-based systems while the top right embedded figure in each sub-figure refers to the Nanopore-based systems. (A) The sequence corruption rate with the assumption of non-consensus method at the receiver. (B) The sequence corruption rate with the assumption of consensus method (i.e., majority selection at each position) at the receiver. (C) The overall sequence error rate consisting of sequence loss and sequence corruption. (D) Theoretical and experimental sequence error rate against channel coverage with the assumption of uneven copy distribution and using non-consensus method. The overall sequence error rate decreases with the increase of coverage.

Next, we generalize the formulation of sequence error rate with uneven copy count distribution. In this case, the sequence error is composed of sequence loss and sequence corruption. The average sequence loss rate 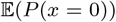 against the average copy count, i.e., the channel coverage (*η*), can be well described by an exponentially decreasing curve *e*^−*λ*^ in which *λ* is a random variable (RV) following an uneven sequence count distribution Λ. The overall sequence error rate against the channel coverage is shown in Fig. 2C, in which the blue and red curves represent the error rate before and after including sequence loss, respectively. Comparing the blue and red curves, we observe that the sequence loss has a more significant impact on the sequence error rate in Illumina-based DNA data storage. On the contrary, the sequence corruption affects the sequence error rate more in Nanopore-based DNA data storage (embedded figure in Fig. 2C).

#### 3.1.2 Elaborated derivation of sequence error rate

We adjust the simplified majority mechanism from binary to quaternary and extend the copy count (the channel coverage) from the constant value *η* to RV *η*_*i*_ subject to certain distribution *H*. Thus, for the non-consensus approach, the expected sequence error rate of Ω_1_ with uneven copy distribution becomes,

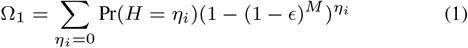

where *η*_*i*_ represents a copy count subject to a distribution *H* (*η*_*i*_ ~ *H*); *ϵ* is the base error rate; and *M* is the sequence length. Fig. 2D compares the sequence error rates of experimental data from (Wang *et al.*, 2019c) with the estimates derived by (1).

With majority selection as the consensus approach, we have Ω_2_,

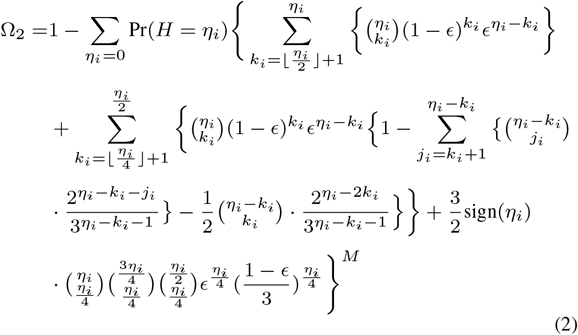

where sign(*x*) is a sign function which equals 1 when *x*(mod 4) = 0 while equals 0 when *x*(mod 4) ≠ 0; and other notations are same as (1). The formulation implies that the biased copy count distribution is not only the origin of the sequence loss but also affects the sequence corruption rate after reconstruction. Specifically, the skewed count distribution of the data at the receiver jeopardizes the overall performance of consensus methods which are usually designed under simplified assumptions, i.e., the distribution of raw data is even or normal. Acknowledging the biased copy count distribution helps better design the consensus algorithms and better estimate the data reconstruction performance from the sequencing data.

#### 3.1.3 Customizing the derivation of sequence error rate with data-dependent errors

In the aforementioned formulations, a constant base error rate *ϵ* is used to represent the probability of random base errors. However, in systems like Nanopore-based systems, some errors occur in a non-random way. These systematic errors (i.e., that are not random) are quantitatively significant and relevant to the features of the reference sequences (or synthesized DNA molecules). For instance, it was found that around 44% of reads of homopolymer runs no less than 5 was observed to contain a deletion error (Lopez *et al.*, 2019). This high error rate and the systematic fashion of the error occurrence aggravate the data recovery difficulty at the receiver. We summarize these errors as data-dependent errors and specify two virtual channels to differentiate the consequences of random and systematic errors (see Supplementary S4). We specifically derive the formulation of the sequence error rate of channels that are prone to data-dependent systematic errors. For the non-consensus approach, we have the expected error rate Ω_3_,

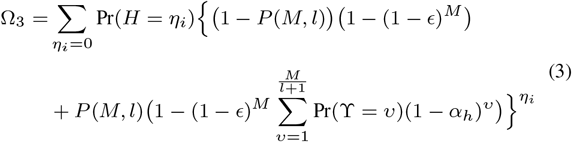

where 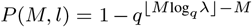 is the probability of a *M*-length *q*-ary sequence having at least one homopolymer larger than length *l* where *λ* is determined by the maximum homopolymer length *l* (see Supplementary S5); 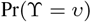 is the probability of a sequence having *v* substrings with homopolymer longer than *l*; *α*_*h*_ is the specific data-dependent systematic error rate; and other notations are the same as (1). For majority selection approach, we have Ω_4_,

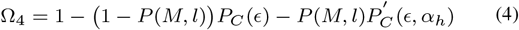

where 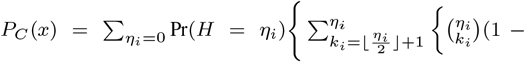 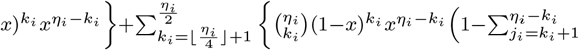 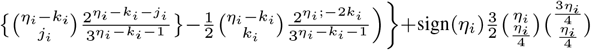 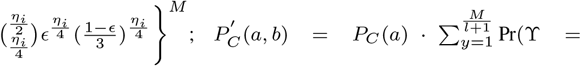 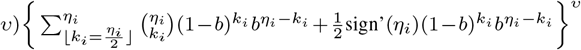 and sign’(*x*) equals 0 when *x* (mod 2) = 1 and equals 1 when *x* (mod 2) = 0. To alleviate the complexities of the formulations, the approximations of (3) and (4) could be found in Supplementary S6. Note that the above formulation is built upon the assumption of storing non-constrained-formatted data where significant increases of systematic errors are present. If data has been constrained-formatted (avoiding the presence of long homopolymer) at the encoding step (Wang *et al.*, 2019a), the severe systematic errors could be avoided in the channel; and the overall sequence error rate of the channel could be measured by (1) and (2) again. However, this constrained formatting/encoding is exploited at the cost of reducing the code rate, i.e., less information stored per nucleotide.

### 3.2 Uneven base-level errors in DNA data storage

#### 3.2.1 Uneven transition errors among nucleotide bases

We first examined the single base error profile. For fair comparison among independent works (Wang *et al.*, 2019b; Heckel *et al.*, 2019; Wang *et al.*, 2019c), we used the filtered reads (i.e., reads with the same length as the encoded sequences) to conduct the analysis. It was found that substitution is the dominant error regardless the choice of sequencing platforms and experimental sets (see Supplementary S7). Furthermore, we analyzed the base transition errors since it could help design more efficient codes as well as to facilitate decoding (Deng *et al.*, 2019). The normalized transition probabilities are shown in Table 1. From the table, we could find that A to C, C to T, T to G, and G to A are the most potential transitions for each reference base, i.e., A, C, T, and G. Regardless of the minor transitions from any of four bases to the N base, the A to C transition is over 3-fold and 4-fold to other two transitions accordingly, i.e., A to T and A to G. Likely, C is almost 3-fold and 4-fold possible to be recognized as T rather than A and G. For T base, the transitions to C and G are both significantly high with approximately 3-fold over T to A transition. Base G has the most considerable transition rates, in which G to A and G to T are about 10-fold than C to G. Overall, G to A, G to T, and A to C are the top three discernible transitions. Several results in the literature are generally consistent with our observations (Pfeiffer *et al.*, 2018; Chen *et al.*, 2014; Ma *et al.*, 2019).

**Table 1.** Base transition error

|  | A | C | T | G | N |
| --- | --- | --- | --- | --- | --- |
| A | - | **0.157** | 0.049 | 0.037 | 0.003 |
| C | 0.032 | - | **0.078** | 0.02 | 0.003 |
| T | 0.021 | 0.062 | - | **0.072** | 0.004 |
| G | **0.201** | 0.073 | *0.182* | - | 0.007 |

The transition error rates among bases, where the row labels denote the transmitting/reference bases and the column labels denote the transmitted bases. The bold values represent the transitions with the highest probability for four reference nucleotides. The italic values represent the overall top 3 erroneous transitions.

#### 3.2.2 Uneven k-mer error patterns

In addition to single base error, we estimated the k-mer error patterns in DNA data storage channel. We first analyzed the 2-mer error patterns with respect to deletions, insertions, and substitutions (Fig. 3A). For deletions, we found that the most erroneous 2-mer patterns of four reference bases are their corresponding 2-mer repetitions i.e., AA, CC, TT, and GG. For insertions, the 2-mer patterns that are prone to having an in-between insertion are AG, CG, TG, and GA for A, C, T, and G, respectively. For substitution error rates of patterns where the first base is substituted to other bases, there is no significant discrimination albeit with the most erroneous 2-mer patterns for each base are the corresponding 2-mer repetitions. The G-oriented patterns show higher substitution rates than the error rates of the other patterns, which concurs with the single base error statistics. Likewise, we analyzed the 3-mer error patterns (Fig. 3B). For deletions, the most error-prone 3-mer patterns for each nucleotide base are the corresponding 3-mer repetitions, i.e., AAA, CCC, TTT, and GGG, which are in line with 2-mer deletion patterns. Moreover, the 3-mer repetitions present higher error rates than the 2-mer repetitions, implying that longer homopolymer might have higher deletion tendency. Again, there are much higher insertion error rates for specific 3-mer patterns, including AGG, CGG, TGG, and GAA. Interestingly, except GCC, all other top 3 erroneous 3-mer patterns (that are shown in Fig. 3B) are with prefixes that are the most error-prone 2-mer patterns, i.e., AG, CG, TG, and GA. This infers that the insertions observed in the received data might highly relate to the neighboring bases. Similarly, most of the top 3 erroneous 3-mer patterns with the first nucleotide being substituted are consistent with the most error-prone 2-mer patterns. However, again, there is no significant discrimination between repetitive patterns and other non-repetitive patterns. Equivalently, the 4-mer repetitions are observed to have the highest tendency toward deletions. And the 4-mer repetitions are with higher deletion rates than the 2-mer and 3-mer repetitions, which further proves that the longer the homopolymer, the higher the probability to encounter deletions (Fig. 3C). For insertions, the most discernible 4-mer patterns for each base are AGGG, CGGG, TGGG, and GAAA. All of them are with 3-mer prefixes that are the most error-prone 3-mer patterns, i.e., AGG, CGG, TGG, and GAA. For most of the 4-mer patterns, we observe similar substitution rates as the 2-mer and 3-mer cases. This suggests that homopolymer does not have significant impact on substitution rates. We also examined errors occurring at the second position in 2/3/4-mer and the last position in 3-mer and 4-mer (see Supplementary S8) and we find that the result is either with no significant erroneous patterns or in line with the patterns observed in the first position.

**Fig. 3.**
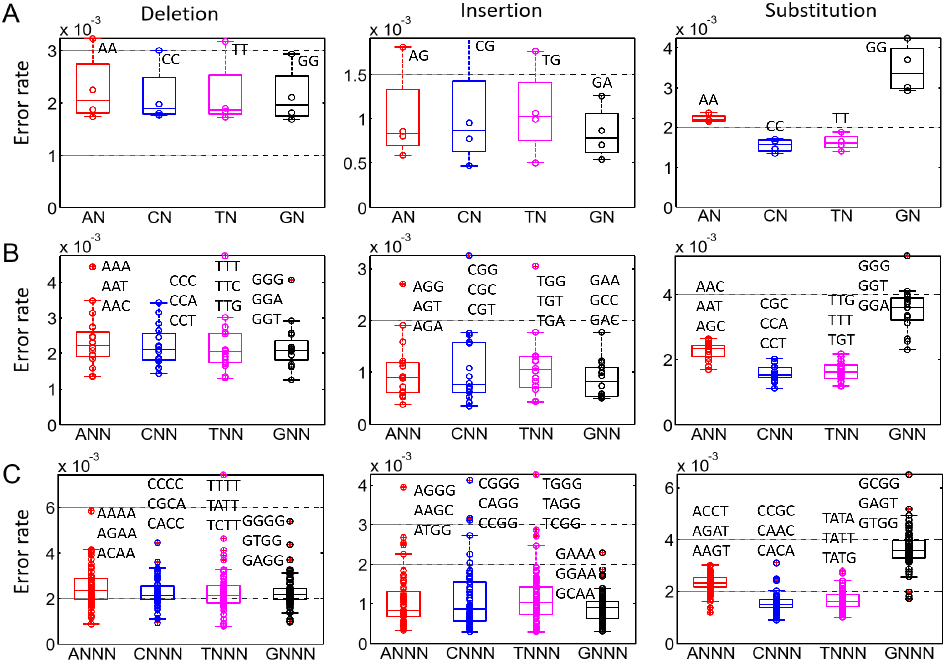
Uneven 2/3/4-mer error patterns for deletions, insertions, and substitutions in DNA data storage, in which the most erroneous patterns are marked correspondingly. With the first base being deleted, following an insertion, and being substituted, the error rates of (A) 2-mer patterns; (B) 3-mer patterns; (C) 4-mer patterns. The homopolymer is observed to have an impact on the deletion errors. There are specific patterns that are prone to have an insertion in-between. No significant discrimination among patterns is observed for substitution errors.

### 3.3 Factors impacting overall sequence error rates in DNA data storage

#### 3.3.1 Sample preparations affect the copy count distribution

To start with, in Fig. 4, we compared the copy count distributions of the reference sequences in our two experiments with different sample preparations (Wang *et al.*, 2019b,c). The scheme in (Wang *et al.*, 2019c) was designed with a single primer binding site (PBS) without conducting PCR amplification before proceeding to DNA sequencing; and the other one (Wang *et al.*, 2019b) was designed with double PBSs and the sample was amplified with 9 cycles of PCR before sequencing. We used the data sets with channel coverage 20x, which means that ideally 20 copy counts of each reference sequence could be found at the receiver. The two observed distributions both approximate to negative binomial distribution which is the consequence of randomly down-sampling from a large population with gamma distribution. Biases in the count distributions are observed for both experiments; and the bias in the single PBS set is larger with size parameter *r* of smaller value, i.e., 2.7 versus 3.3. Besides, in the single PBS set, one reference sequence has 174 copy counts far away from the mean coverage 20x. The different degrees of PCR bias and PCR stochasticity (see Supplementary S9) might ascribe to the bias difference between the two experiment sets. To further confirm the major source of the bias differentiation between two distributions, we compared the copy count distributions of all sequences, sequences including 4nt homopolymer, and sequences without 4nt homopolyer. It is observed that the sequence with maximum copy counts, i.e., 174, is with 4nt homopolymer. And there is no obvious discrepancy in distributions among three sets (see Supplementary S9 Fig. 14). This implies that rather than distinct PCR biases caused by sequence-specific randomness (due to homopolymer differentiation), distinct degrees of PCR stochasticity caused by different sample preparations majorly explain the bias difference.

**Fig. 4.**
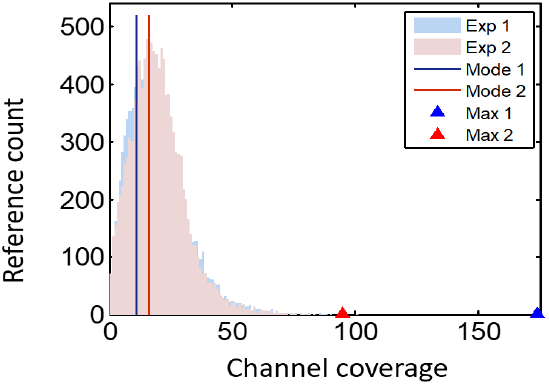
Copy count distributions for varied sample preparations. Based on data sets with 20x channel coverage, the blue-colored copy count distribution is from the single primer binding site (PBS) experimental set; while the red-colored copy count distribution is from double PBS experimental set. The blue and red solid lines represent the modes of two distributions, respectively. The maximum counts observed in two sets are triangle marked, and there is one reference sequence with 174 copy counts in the single PBS set (i.e., blue triangle). Both distributions approximate to negative binomial distribution, the single PBS distribution is with higher bias where the size parameter *r* is smaller. i.e., 2.7 versus 3.3.

#### 3.3.2 Sequence structure and downstream processing affect the base error, sequence corruption, and system capacity

The basic data unit in DNA data storage, i.e., DNA sequence, is usually designed with length ~ 200 to tailor the current synthesis and sequencing techniques. For Illumina-based systems, a PE150 protocol could be used to increase the sequencing accuracy by stitching two paired-end (PE) reads. We aligned the merged reads back to the reference sequences to study how the reference sequence length and the general merging processing cooperatively affect the error profile in DNA data storage channel. The average positional error rates along the coordinate of the reference sequence are shown in Fig. 5. The average base error rate along the coordinate is uneven where the overlapped region (pink-colored in Fig. 5) has lower error rate than the non-overlapped region (blue-colored) does. And the substitution errors are reduced most notably in the overlapped region. The overlapped region is a region corresponds to the universal regions shared by two PE reads. The unevenness between overlapped and non-overlapped regions is due to the gap between the length of reference sequences (i.e., 190nt and 190~199nt) and the length of PE reads (i.e., 150nt). Moreover, by comparing the blue-colored (i.e., non-overlapped) regions in Fig. 5A, we could find that the non-overlapped region with a PBS region (gray-colored) as the adjacent has a lower error rate than the other non-overlapped region.

**Fig. 5.**
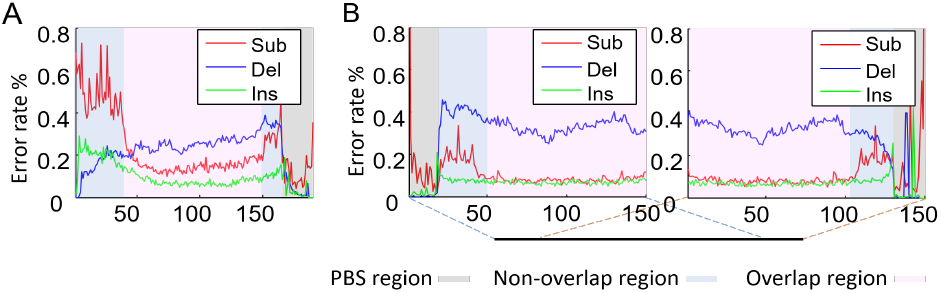
Positional error profile of reference sequence in DNA data storage using PE150 sequencing protocol and merging processing, in which three regions are highlighted along the coordinate. The positional error rate profile of (A) reference sequences with length of 190nt. (B) reference sequences with lengths ranging from 190nt to 199nt. Two 150-length positional error profiles starting from two ends of the reference sequences are presented simultaneously to accommodate the variable-length feature of the reference sequences.

Additionally, we analyzed the data sets that have been filtered by lengths. This further proves structural design of the sequence, i.e, appended with single PBS or double PBS and fixed length or variable lengths, affects the data integrity at the decoder (see Supplementary S10). Despite that filtering alleviates errors especially indels, filtering might aggravate sequence loss especially when the number of reads provided at the sequencer is limited. Hence, we compared the overall sequence rates before and after filtering with given amounts of reads (see Supplementary S11). By incorporating the sequence loss and sequence corruption (under the trial-and-error assumption), the filtered data set shows higher overall sequence error rates than those of non-filtered data set up to coverage ~ 30x (Supplementary S11 Fig. 16C). This suggests that while filtering might immediately reject seemingly erroneous data, the sequence loss caused by it might be more significantly increase the overall sequence error rate, rendering more stringent requirements on the code design.

Inspired by the unevenness of base error rates along the sequence coordinate caused by the length difference between reference sequences and PE reads, we theoretically analyzed the impact of the sequence length on the sequence corruption rate. First, Fig. 6A depicts how the ratio of read length to the sequence length affect the sequence corruption rate. Specifically, when the ratio is below 0.5, stitching two PE reads is unable to recover the reference sequence (see Supplementary S1 Fig. 1C), leading to 100% corruption. When the ratio is from 0.5 to 1, the corruption rate decreases with the increase of the ratio. When the ratio is no less than 1, each PE read could ideally cover the whole sequence (see Supplementary S1 Fig. 1A), rendering the merged read with a fully overlapped region and consequently reducing the corruption rate to a low floor. Comparing the curves with different colors, the impact of the ratio on the sequence corruption rate was found more noticeable for shorter sequence length. With PE read length 150, Figs. 6B and C show that the sequence length directly affects the sequence corruption rate and incremental redundancy for addressing the corruption.

**Fig. 6.**
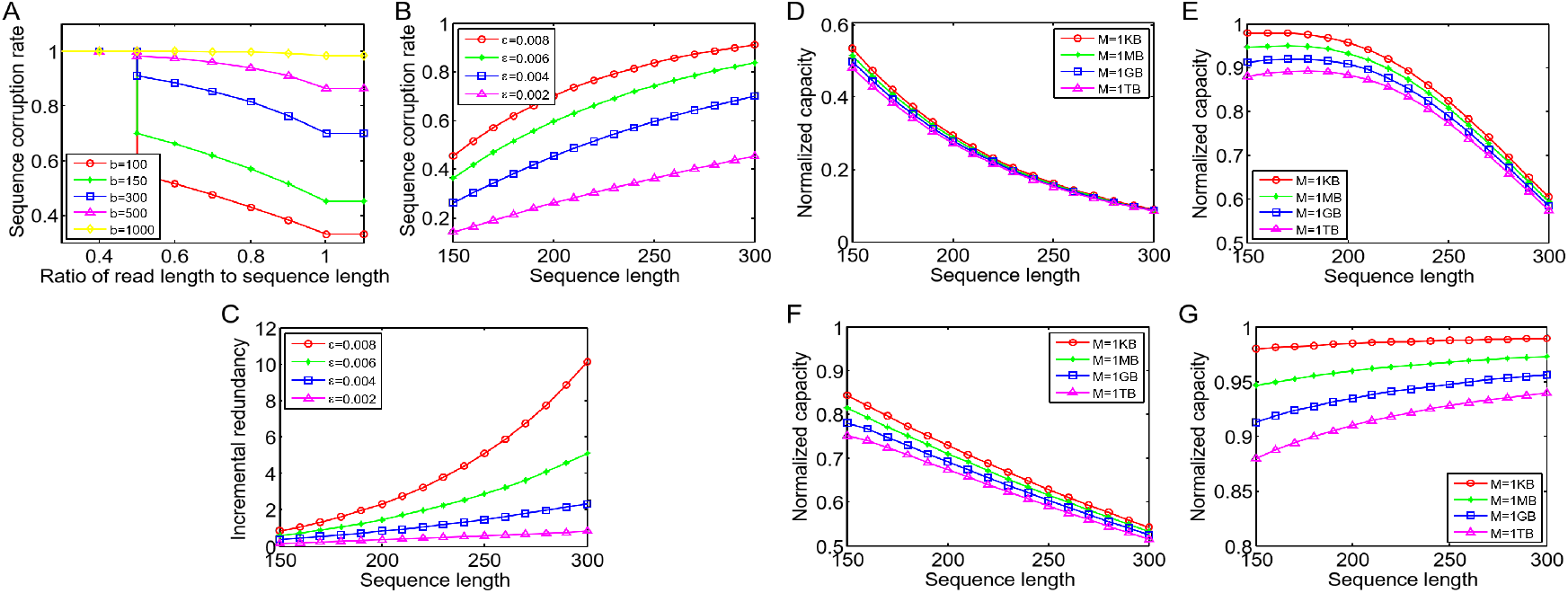
Theoretical analysis of the impact of sequence length. (A) Sequence corruption rate against the ratio of PE read length to sequence length. Curves with different colors represent different sequence lengths *b*. With PE read length 150 and curves with different colors representing different base error rates *ϵ*, (B) sequence corruption rate against sequence length; (C) incremental redundancy for addressing sequence corruption against sequence length. The achieved capacity (normalized) against the sequence length, (D) when channel coverage *η* = 1 and raw base error rate *ϵ* = 0.8%; (E) when *η* = 10 and *ϵ* = 0.8%; (F) when *η* = 1 and *ϵ* = 0.2%; (G) when *η* = 10 and *ϵ* = 0.2%. Curves with different colors represent different sizes of stored data *M*.

We continue to explore the impact of the sequence length on the achieved capacity where the capacity is determined by the redundancy required for correcting the sequence corruption and the redundancy required for indexing. These two redundancies are highly related to the sequence length; and the impacts of sequence length on them are opposite. With different channel coverage *η* and base error rates *ϵ*, Figs. 6D-G show how the achieved capacity change with the sequence length in which different colors represent different stored data sizes ranging from 1 Kilobyte to 1 Terabyte. We found that for higher base error rate systems (Figs. 6D and E), the achieved capacity decreases with the increase of sequence length, presenting that the impact of increased redundancy for error correction on the capacity plays the prime role. This trend also appears in lower base error rate systems when the coverage is 1x (Fig. 6F), i.e, ideally only one read for each reference sequence could be used for data reconstruction. Interestingly, the trend reverses in the lower base error rate system when the coverage is 10x (Fig. 6G). The much lower corruption rates ascribed to the high coverage could be the reason of the reversed trend. The error correction redundancy required by the much lower corruption rates no longer weighs higher than the indexing redundancy in regard to affecting the capacity. Therefore, in Fig. 6G, the increased capacity with the increased sequence length is mainly due to the decreased indexing redundancy. This indicates that in most cases, designing sequences with short length could improve the achieved capacity (Figs. 6D-F). However, for storage systems with very low raw base errors and sufficient coverage at the receiver (Fig. 6G), sequence length could be designed as long as possible (up to twice of PE read length) to improve the achieved capacity.

#### 3.3.3 Theoretical estimation of the overall sequence error rate

Using different synthesis techniques and experimental settings gives different count distributions at the receiver, leading to distinct sequence loss rates. Meanwhile, using different sequencing techniques and data processing methods gives different base error rates at the receiver, leading to distinct sequence corruption rates. We thus theoretically estimate the overall sequence error rate consisting of sequence loss and sequence corruption by setting a range of practical values to several important impacting factors. With the assumption of using non-consensus post-processing method after merging PE reads, the formulation of the overall sequence error rate is an extended version of 1 where the base error is no longer constant along the coordinate but varies with the region, i.e., overlapped or non-overlapped.

We separately consider two scenarios. First, the merged reads are with fully overlapped region. To comply with the assumption, the sequence length is set equal to the PE read length (i.e., 150nt). The sequence error rate against the channel coverage is illustrated (see Supplementary S13). In general, the higher the base error rate is, the higher the channel coverage is required to achieve a similar sequence error rate at the decoder. The variations among different base error rates, i.e., among sub-figures, are not notable. In the aspect of copy count distribution, the smaller the size parameter is, i.e., the more over-dispersion of the distribution, the higher the sequence error rate. Next, we analyzed the non-overlapped read case by setting the sequence length twice of the PE read length. Similarly, we draw two sub-figures, i.e., Figs. 7A and B, corresponding to two different raw base error rates, i.e., 0.8%, 0.3%, corresponding to Miseq and Hiseq Illumina sequencing, respectively (Quail *et al.*, 2012). The trend of the sequence error rate against the channel coverage in this case is the same as the overlapped case, but the discrimination among different base error rates, i.e., among sub-figures, is more notable. Comparing these two groups of figures (Fig. 7 versus Supplementary S13 Fig. 17), it could be found that to obtain similar sequence error rate with the same base error rate and size parameter, the required coverage of the non-overlapped case is no less than the overlapped case, i.e., ~ 4-fold for 0.8% base error rate, ~ 1.5-fold for 0.3%. To this end, we conclude that if the raw base error rate could be kept around 0.3% or less, the sequences could be designed with long length (i.e., from 150 to 300) where merged reads are all non-overlapped. However, for systems with higher base error rate, short sequence design (i.e., no longer than 150) which leads to all merged reads overlapped or semi-overlapped is a better choice.

**Fig. 7.**
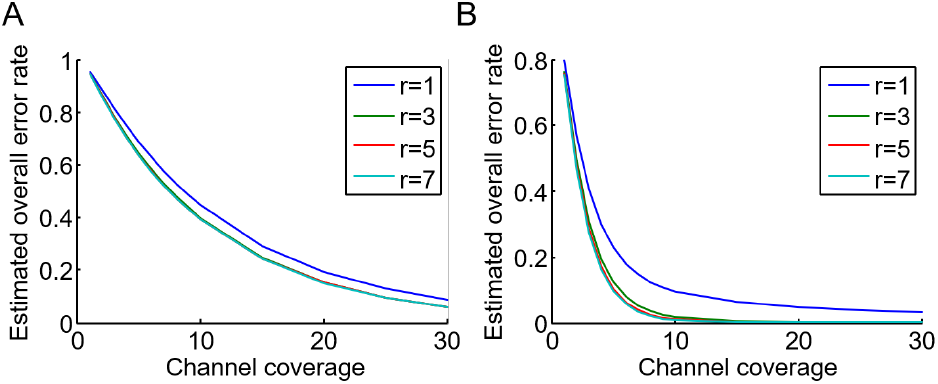
With the assumption that the sequence length is twice as the read length where the merged reads are with the full non-overlapped region, the overall sequence error rate in Illumina-based storage against the channel coverage, where the raw base error rate is (A) 0.8% similar to Miseq; (B) 0.3% similar to Hiseq (Quail et al., 2012). In each figure, different colors represent different size parameters of the copy count distribution that are ascribed to sequence loss.

## 4 Discussion

Distinct from other traditional storage systems, DNA data storage systems exhibit few unique characteristics. Specifically, there are generally amount of redundant data copies of original data albeit these copies might be corrupted by base errors; and the number of the redundant data copies for each original data unit are uneven. Most of existing works (Erlich and Zielinski, 2017; Heckel *et al.*, 2019) only discussed the adverse consequence of the uneven copy count distribution, i.e., sequence loss, while overlooked the multi copies’ benefit to data reconstruction. Also, physical redundancy of data copies at the decoder was excluded from the channel and discussed separately from the logical redundancy of error control code. In fact, the multi-count data feature of DNA data storage enables a pre-decoder data reconstruction from multi (erroneous) copies where the failure of the reconstruction is termed as sequence corruption. With the preliminary reconstruction before decoding, the data imperfection at the decoder that consists of sequence loss and sequence corruption offers a unified error profile for DNA data storage channel, easing the channel analysis and giving new insights for future code design.

Diving into the data imperfection observed from the experiments, biases have been found both in copy counts and base error patterns. The existence of these biases further distinguishes the DNA data storage from other conventional storage systems, suggesting that facilitating higher performance gains in terms of capacity, reliability, and robustness in DNA data storage are possible. Moreover, the unevenness in the error rates of base patterns, including uneven error rates of single base transition and k-mer deletion/insertion, could be used as prior knowledge for decoding and optimizing encoding. For instance, using the transition tendency as the additional information to the decoder increases the error correction performance (Deng *et al.*, 2019). Also, the uneven transition feature could be leveraged to design unequal codes with higher efficiency. In addition, the deletion-prone characteristic in the long homopolymer patterns especially in Nanopore-based systems suggests that coding techniques that restrict the homopolymer length, i.e., constrained coding (Wang *et al.*, 2019a; Immink and Cai, 2017; Song *et al.*, 2018), might be a promising solution provided that the subsequent reduction on code rate/capacity is tolerable. Additionally, with the prevalent PE sequencing protocol and merging processing as the premises, the uneven error rate along the sequence coordinate (between non-overlapped and overlapped regions) is another unevenness that could be used for code design, e.g., unequal encoding, to increase the achieved capacity.

In this work, the impact of sample preparation on the data at the receiver was investigated based on two experiments with different rounds of PCR amplifications before DNA sequencing. Specifically, the amount of the copy count before sending to sequencing which is the consequence of the sample preparation (i.e., PCR) affects the bias observed in the sequenced data. And this bias subsequently causes sequence loss that pertains to the sequence error threatening on the decoder. We have shown that samples with more rounds of PCR attribute to less biased sequenced data since it provides more sufficient initial amount of molecules which avoids severe PCR stochasticity. Besides PCR, other sample preparation steps should also affect the data integrity at the receiver and could be investigated in the future work. The impact of sequence length on the data imperfection at the decoder and channel capacity was also theoretically studied. It was observed that only with sufficient coverage and low base error rate, the achieved capacity could increase with the increase of sequence length. However, in other cases, the achieved capacity decreases with the increase of the length because of the increased redundancy for addressing increased corruption rate. Hence, the system should be designed through comprehensive consideration of the involved impacting factors and the trade-offs among them.

## 5 Conclusion

In this work, we have conducted a comprehensive investigation of errors in DNA data storage channel. Quantitatively, the data imperfection including sequence loss and sequence corruption at the decoder has been presented. Besides deriving the sequence error rate to monitor the data reconstruction demand, we also further studied on the imperfect data and found out that unevenness exists in several aspects and it could in turn to help designing systems with better performance. Additionally, we experimentally and theoretically analyzed the sequence error rates under different experiment settings and various but realistic parameter settings, including sequence lengths, base error rates, and over-dispersion degrees of distribution. From the perspective of data reconstruction, the results reported provide new perspectives for the development of the advanced future DNA data storage.

## Acknowledgements

We would like to thank Sergio Peisajovich and Saurabh Nirantar from Illumina, and Luo Lei from NovogeneAIT. We thank Ng Yi Mei for assisting on the initial analysis.

