## Supplementary for "Modelling, characterization of data-dependent and process-dependent errors in DNA data storage"

### S1. Pair-end (PE) sequencing protocol and merging post-processing technique

With the pair-end (PE) sequencing protocol, for one oligo molecule, Illumina sequencer can generate a pair of read sequences starting from two ends of the oligo. One of the most significant advantages of PE protocol is merging the two reads could increase the accuracy of reading the sequenced molecule. As illustrated in **Supplementary S1 Fig. 1A**, there is a pair of forward and reverse reads corresponding to one original DNA oligo. If the length of the original oligo is as short as the read's length, each read can cover the whole oligo while if the oligo is longer than the reads but shorter than twice the read, i.e., **Supplementary S1 Fig. 1B**, only two PE reads stitching together could potentially recover the original oligo. However, if the oligo is longer than twice the PE read, i.e., **Supplementary S1 Fig. 1C**, combining two PE reads still could not recover the original oligo which partially upper bounds the longest DNA sequence that could be read with ensured integrity. Most existing works (1–6) have used PE150 sequencing protocol, i.e., 150nt reads from two ends of the original molecule, for reading data in DNA storage. As a widely adopted post-processing method for sequencing data, the generated PE reads are usually stitched together through widely recognized computational tools, e.g. (7, 8), before proceeding to further analysis. However, these tools could not guarantee every pair of PE could be successfully merged into an assembled sequence and some merged data are with notable errors, i.e., with lengths differ from the oligo's length. To keep consistency with (4), the PE150 reads generated in our two previous works (5, 6) were merged using FLASH (8) with parameters setting -m 10 -M 100 -x 0.1 -p 33, where -m refers to minimum overlap length; -M refers to maximum overlap length; -x refers to the maximum allowed ratio of mismatch to the overlap length; and –p refers to the acceptable quality score of PE reads. One instance resting on (5) is shown in **Supplementary S1 Fig. 2**, where the merged reads are with lengths ranging from 150nt to 290nt, varied from the original oligo's length, i.e., 190nt. The range of the length relates to the parameter setting and might be varied from case to case. The instinct following-up processing is filtering based on the length while it might pose an increase in the sequence loss which is detrimental to data reconstruction. To that end, the processing method on the merged reads should be decided after considering all trade-offs.

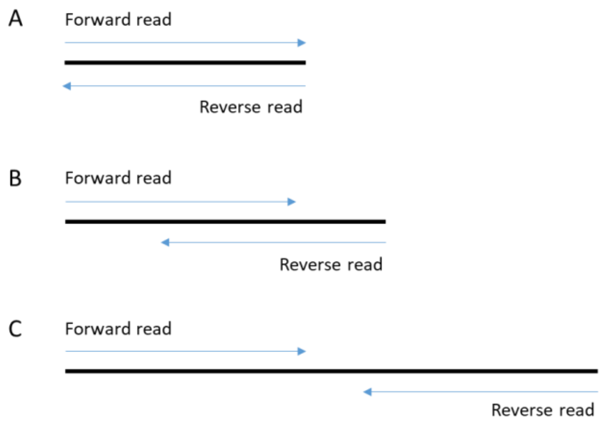


**Supplementary S1 Fig. 1 Three scenarios of the generation of PE reads for the original oligo.**

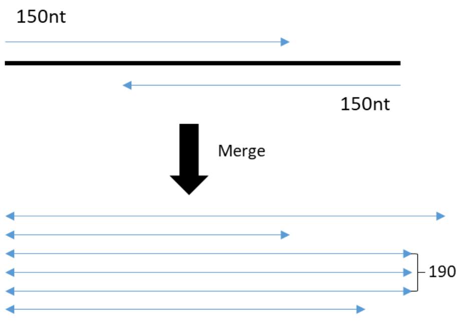


**Supplementary S1 Fig. 2 The potential results of merging PE reads.**

#### S2. Simulation and evaluation of the uneven multi-count distribution in a computational model

The sequence loss/dropout that constitutes the sequence error stems from the uneven multi-count distribution phenomenon where certain original sequences have no copy being observed at the receiver. More precisely, the sequence loss could be further categorized into physical molecule loss and sequencing data loss. And these two losses are relevant to the multi-count distribution at specific stages, i.e., storage and sequencing. To trace the change of the multi-count distribution and predict the sequence loss that might threaten the decoder, a computational model has been proposed in (9). By exploiting the model, we elaborate on the impacts of the two mentioned sequence losses on the resultant loss. With a variety of storage density ($\eta_{store}$) that relate to physical molecule loss and sequencing coverage ($\eta_{seq}$) (which is called channel coverage in the main text) that relate to sequencing data loss, the physical sequence loss $\delta_1$ and overall sequence loss $\delta_{all}$ against sequencing coverage are depicted in **Supplementary S2 Fig. 3**. It could be found that the overall sequence loss decreases with the increase of sequencing coverage in an exponential way consistent with the Poisson sampling effect. On the other hand, when the sequencing coverage is sufficient, the overall dropout rate is lower bounded by the physical molecule loss that rests on the storage density. Moreover, to figure out how the multi-count distribution of synthesized molecules impacts the sequence loss, with different colours representing distributions with different biases, i.e., coefficient of variations (CVs), the sequence loss/dropout rate against the storage coverage (plotted with logarithm) is shown in **Supplementary S2 Fig. 4**. The illustration shows that with multi-count distribution with higher dispersion, i.e., smaller C.V., there is severer sequence loss, suggesting that controlling the dispersion of the multi-count distribution of molecules at the synthesis stage could suppress the sequence loss at the storage stage. In addition, the association of the overall sequence loss ($\delta_{all}$) and the physical molecule loss ($\delta_1$) is drawn in **Supplementary S2 Fig. 5**. From the figure, it could be noticed that when $\eta_{seq}$ is insufficient, e.g., less than 10x, the $\delta_{all}$ changes non-linearly with the $\delta_1$ especially when $\delta_1$ is with insignificant values. This further proves that albeit the overall sequence loss is lower bound by the physical molecule loss, the scarcity of sequencing coverage's impact on the sequence loss should not be underestimated.

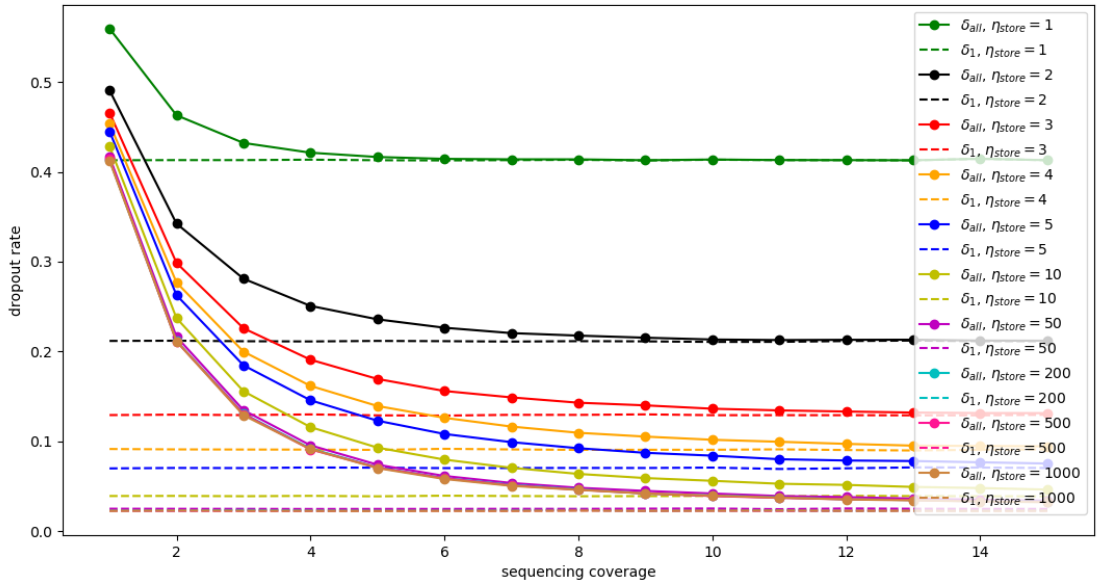


**Supplementary S2 Fig. 3 The sequence loss/dropout rate against the sequencing/channel coverage, in which the dotted lines represent the physical sequence loss $\delta_1$ and the solid lines represent the overall sequence loss $\delta_{all}$ composed of physical sequence loss and sequencing loss. Each pair of lines with distinct colours corresponds to a given storage density $\eta_{store}$.**

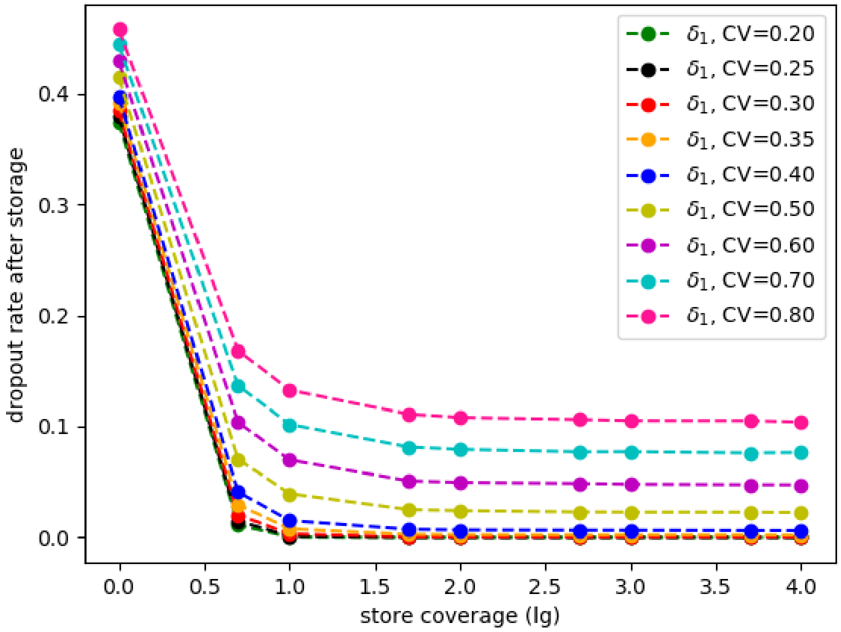


**Supplementary S2 Fig. 4 The sequence loss/dropout rate against the storage coverage (with logarithm), in which curves with different colours refer to multi-count distributions with different coefficient of variations (CVs) that represent different dispersions.**

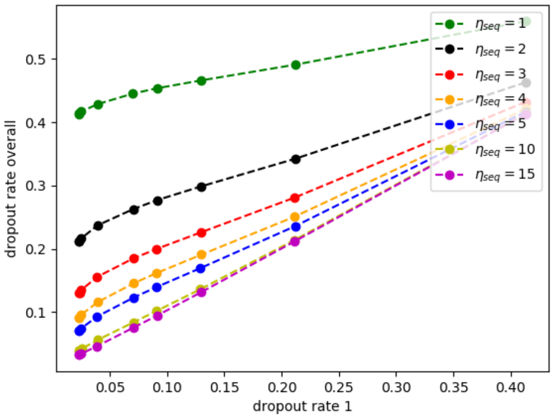


**Supplementary S2 Fig. 5 The overall sequence loss/dropout rate ($\delta_{all}$) against the physical molecule loss ($\delta_1$) with different curves representing different sequencing/channel coverage ($\eta_{seq}$)**

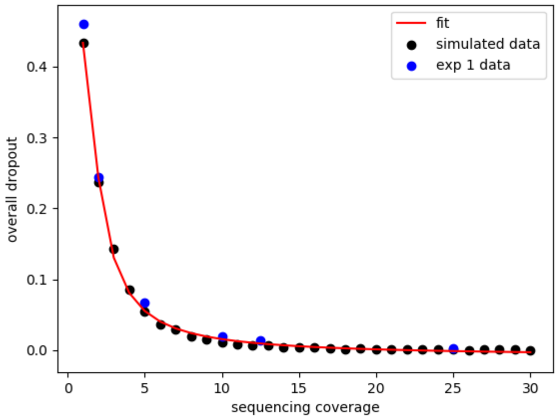


**Supplementary S2 Fig. 6 The overall sequence loss/dropout rate ($\delta_{all}$) against the sequencing coverage/channel ($\eta_{seq}$) where the black dots are the simulated data with the red line as fitting while the blue dots are data from the experiment.**

Furthermore, we evaluate the computational model in terms of predicting the overall sequence loss with different sequencing coverage. To do that, the simulated result from the model is compared with the result using the data collected from our previous work (5). As shown in **Supplementary S2 Fig. 6**, the sequencing sampling effect simulated by this model is well-fitted with the experimental data with $R^2 = 0.96$.

In addition to the sequencing process, the effect of the storage process on the sequence loss simulated by the model is further evaluated by comparing with the dilution experimental results in (1). (1) has reported sequence loss results of reading data from sets of diluted samples, i.e., from 10fg~10ng, which correspondingly refer to storage densities from 1 copy per sequence to $10^6$ copies per sequence. Accordingly, the simulated sequence loss and multi-count distribution are generated by providing a serial of values of storage density modelling the dilution effect on the modified model. Instead of direct dilution used in the original model in (9), we modified the model to simulate the serial dilution by continuously iterating the sampling process until a specific dilution level is achieved. This minor modification could better simulate the dilution experiments used in (1). However, with other parameters setting similar to (1), i.e., 40 PCR cycles for sample preparation, and ~400x sequencing coverage, notable gaps between the simulated results and experimental results are observed. Specifically, for storage densities 10pg ($10^3$ physical copies/seq), 1pg ($10^2$ physical copies/seq), 100fg (10 physical copies/seq), and 10fg (1 physical copy/seq) , there are still notable gaps between the experimental results and the simulated results, i.e., 4%, 62%, 80%, and 87% versus 0.9%, 0.92%, 1.69%, and 41.63% (see **Supplementary S2 Fig. 7**). The losses are predominated by the physical molecule loss at the storage stage, i.e., the sequence losses after storage with given densities are 0.28%, 0.31%, 0.92%, and 41.45%. This is because that while PCR amplification following storage causes severe skewness on the multi-count distribution owing to the PCR bias or PCR stochasticity, the sequencing depth in the subsequent sequencing process is high enough to overcome the Poisson sampling effect, alleviating the additional sequence loss at the sequencing sampling stage (10).

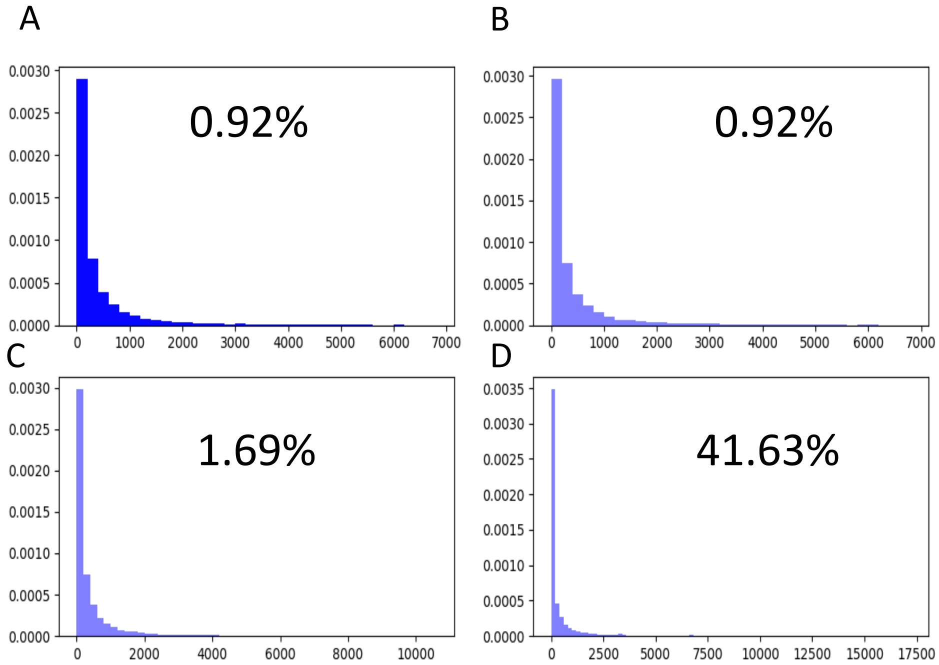


**Supplementary S2 Fig. 7 The simulated multi-count distribution by the computational model with storage densities of A. 10pg (1000 physical copies/seq), B. 1pg (100 physical copies/seq), C. 100fg (10 physical copies/seq), and D. 10fg (1 physical copy/seq) when the sequencing/channel coverage is 400x. The percentages in each sub-figure are the sequence loss rates.**

Note that, the simulated results presented above are based on setting the PCR amplification efficiency as a random variable (RV) subject to a normal distribution, which is a widely accepted assumption. Furthermore, to figure out if the observed gap between the experimental results and the simulated results are relevant to the used PCR amplification efficiency, without changing other parameter settings, we test the model by using two other types of value for PCR efficiency, i.e., constant (**Supplementary S2 Fig. 8A**) and strand-specific RV (**Supplementary S2 Fig. 8C**). Besides the increasing sequence loss and dispersion with the increase of PCR cycles, it could also be found that both the sequence loss and dispersion of the multi-count distribution are severer with the assumption of the strand-specific RV (**Supplementary S2 Fig. 8C**) due to the PCR bias, approximating the sequence loss observed in the experimental data in (1).

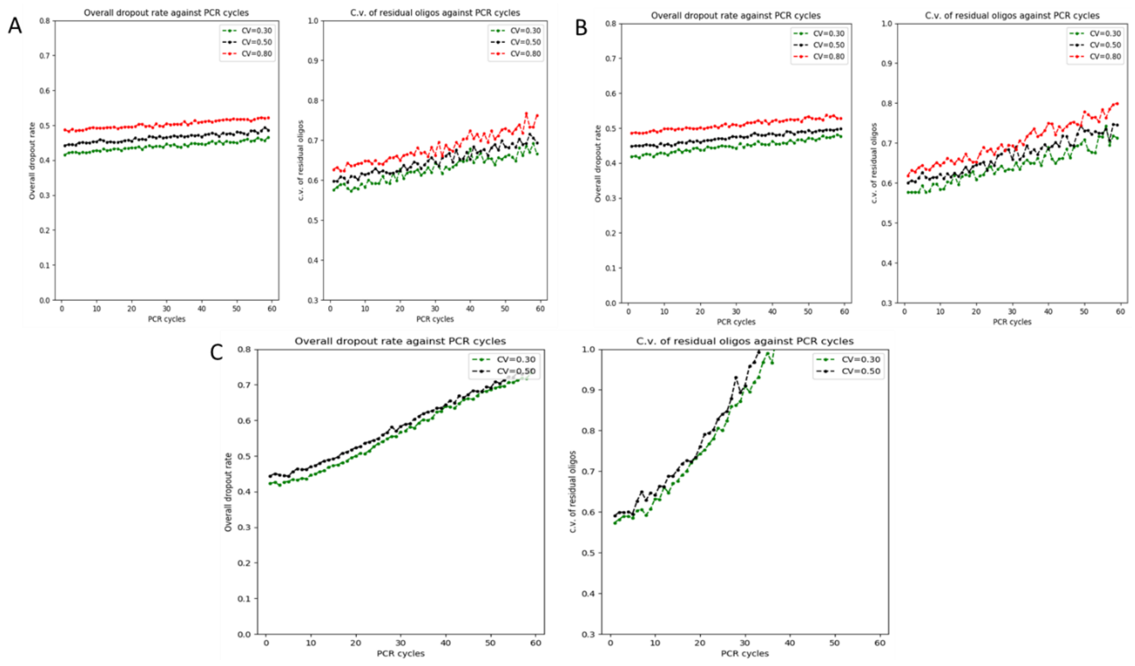


**Supplementary S2 Fig. 8 The simulated sequence loss rate and coefficient of variation (C.V.) of the multi-count distribution when PCR efficiency is A. a constant value $\mathbf{p = 0.85}$ for each cycle pf each strand; B. a random variable (RV) $\mathbf{p \in \mathbb{N}(u = 0.85,\ \sigma = 0.07)}$ for each cycle of each strand where the RV follows a normal distribution with mean value 0.85 and standard deviation 0.07; C. a strand-specific RV $\mathbf{p \in \mathbb{N}(u = 0.85,\ \sigma = 0.07)}$ for each strand, and the value is fixed for each cycle. The curves with different colours in each sub-figure represent results from samples with different initial distributions, i.e., different C.V..**

### S3. The customized parameters for monitoring the sequence error rate

Based on the practical measures, the base error rate $\varepsilon$ is set to 0.1%, and the sequence length $M$ is set to 200 to simulate the sequence corruption of Illumina-based DNA storage, i.e., $\Omega_{illu}(\eta; \varepsilon, M)$. For simulating the Nanopore-based DNA storage, i.e., $\Omega_{illu}(\eta; \varepsilon, M)$, the base error rate $\varepsilon$ is set to 10% and the sequence length $M$ is set to 1000. For the trial and error (non-consensus) case, i.e., the sequence is considered as corrupted only if all copies of the sequence are erroneous, the curves representing the association between sequencing/channel coverage and sequence corruption (Figure 2A in main text) are based on formula $\Omega = (1 - (1 - \varepsilon\,)^{M})^{\eta}$. When the sequencing/channel coverage is 1x, i.e., for each original sequence, there is only one copy could be used for reconstruction at the receiver. The same parameter values are used for drawing Figure 2B which is the consensus case with simplified majority selection at each position.

### S4. Multi-count and data-dependent system channel models

It has been reported that in Nanopore-based DNA storage, sequences with specific patterns, i.e., long homopolymer, tend to encounter systematic errors with high probability (9). To separate the distinct demands that originate from the different data formats and different sequencing techniques that have been used for data reading, two virtual data-dependent channels are used to illustrate the discrepancy in **Supplementary S4 Fig. 9**. Specifically, data without constraining the homopolymer might be corrupted by additional deletions at the homopolymer positions. Note that the specific position with one deletion is trivial to the homopolymer as the homopolymer pattern consists of repetitive bases. Compared with the random base errors that stochastically occur in stochastic positions of the sequence, the homopolymer-related errors show an extremely higher error rate and occur systematically, i.e., a large part of the read copies of the sequence is corrupted by the deletion errors at the homopolymer coordinates. Instinctively, this systematic error could not be effectively resolved by a consensus algorithm which is with more proficiency in correcting random errors. Moreover, considering that the sequence corruption rate closely relies on the base error rate, the notably increased error rate in the homopolymer-related deletions is deemed to lead to a significant difference from the random base error. Due to these reasons, it is of critical importance to incorporate the effect of this systematic and highly possible errors into the derivation of the sequence corruption that constitutes the reconstruction demand of DNA storage.

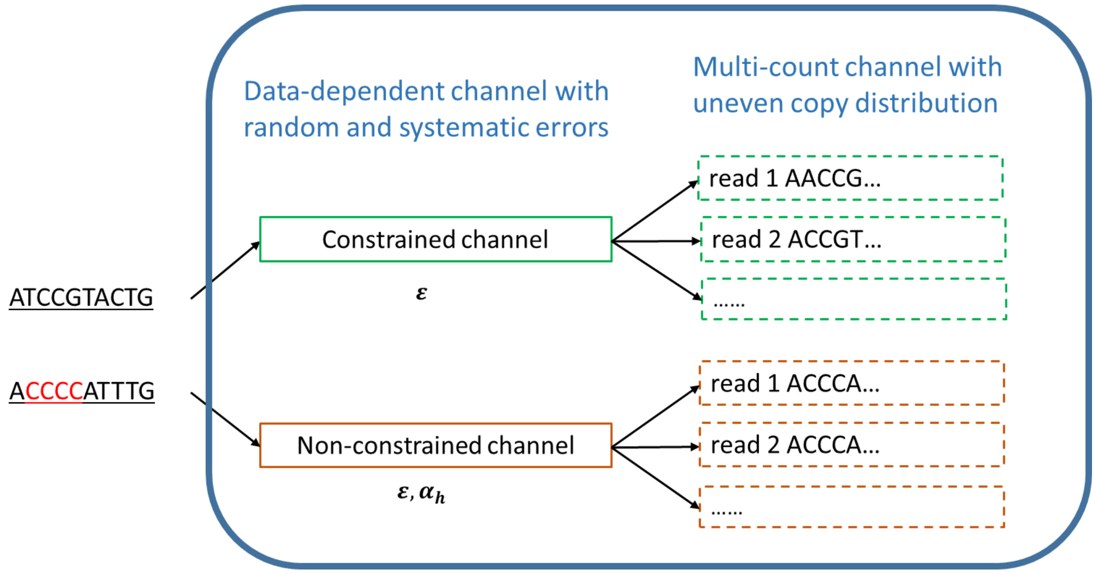


**Supplementary S4 Fig. 9 Two virtual channels for multi-count and data-dependent system. In the constrained channel, sequences without homopolymer, e.g., ATCCGTACTG, are corrupted by random base errors with rate $\varepsilon$ while in the non-constrained channel, sequences that are with homopolymer, e.g., ACCCCATTTG, are corrupted by systematic error $\alpha_h$ that relates to homopolymer in addition to random base error $\varepsilon$.**

#### S5. The capacity of constrained systems

With the constraint of the maximum homopolymer on the encoded DNA sequences, the DNA storage system could be considered as a constrained system. In the aspect of coding, the constrained system is usually considered when the channel presents different characteristics for different data. To suppress the occurrence of errors that relate to the certain data format, constrained codes are leveraged. However, the alleviated errors brought by the constrained codes come with the sacrifice of the reduced capacity or so-called code rate. A homopolymer-limited system like DNA storage can be denoted by $(M, d+1, k+1)$ where $d+1$ and $k+1$ are the lower and upper boundaries of the allowable numbers of continuously repetitive $M$-ary symbols. The system can also be represented by an $(M, d, k)$ finitestate transition diagram (FSTD). The capacity of a constrained system could be derived from the FSTD. The FSTD has $k+1$ states and a $(k+1)*(k+1)$ adjacency matrix $D(dij)$, where each entry $dij$ is the number of edges in the FSTD transiting state $i$ to state $j$. The Markov model can also be used to derive the Shannon capacity of the constrained system using $C = \log_2 \lambda$, where $\lambda$ is the largest eigenvector of $D(dij)$. For constrained DNA storage, there is only one constraint for the maximum homopolymer, so that the system can be denoted as $(M, 1, l)$ where $l$ is the maximum allowable homopolymer. Examples of constrained DNA storage including the FSTD illustration and capacity generation could be easily found in (6, 11).

### S6. The approximation of sequence error rate formulations

With the assumption of using trial and error method as post-processing, the sequence error rate formulation of systems with biased multi-copy distribution and data-dependent errors can be approximated to $\sum_{\eta_i=0} \Pr(\Pi = \eta_i)\left\{\left(1 - P(M,\, l)\right)\left(1 - (1 - \boldsymbol{\epsilon})^{M}\right) + P(M,\, l)\alpha_h\right\}^{\eta_i}$, where $P(M,\, l) = 1 - q^{\lfloor M \log_q \lambda \rfloor - M}$ is the probability of $M$-length sequence having at least one homopolymer larger than length $l$; and of which $q = 4$; and $\lambda$ is dependent on the maximum homopolymer constraint. $\Pr(\Pi = \eta_i)$ is the probability that the sequence has $\eta_i$ copies where the copy number is subject to a distribution $\Pi$. $\boldsymbol{\epsilon}$ is the general base error rate and $\alpha_h$ is the homopolymer-related error rate. On the other hand, with the assumption of using majority selection as post-processing, the sequence error rate formulation of systems with biased multi-copy distribution and data-dependent errors can be approximated to $1 - \left(1 - P(M,l)\right)Pc(\boldsymbol{\epsilon}) - P(M,l)\, Pc(\check{\boldsymbol{\epsilon}})$ , where $\check{\epsilon} = 1 - \sqrt[M]{1 - \alpha_h}$ and $Pc(\mathrm{x}) = \sum_{\eta_i=0} \Pr\,(\Pi = \eta_i)\left\{\sum_{k_i=\lfloor \frac{\eta_i}{2} \rfloor+1}^{\eta_i} \left\{\binom{\eta_i}{k_i}(1-\mathrm{x})^{k_i}\mathrm{x}^{\eta_i-k_i}\right\} + \sum_{k_i=\lfloor \frac{\eta_i}{4} \rfloor+1}^{\frac{\eta_i}{2}} \left\{\binom{\eta_i}{k_i}(1-\mathrm{x})^{k_i}\mathrm{x}^{\eta_i-k_i}(1 - \sum_{j_i=k_i+1}^{\eta_i-k_i}\left\{\binom{\eta_i-k_i}{j_i}\frac{2^{\eta_i-k_i-j_i}}{3^{\eta_i-k_i-1}}\right\} - \frac{1}{2}\binom{\eta_i-k_i}{k_i}\frac{2^{\eta_i-2k_i}}{3^{\eta_i-k_i-1}})\right\} + \mathrm{sign}(\eta_i)\frac{3}{2}\binom{\eta_i}{\frac{\eta_i}{4}}\binom{\frac{3\eta_i}{4}}{\frac{\eta_i}{4}}\binom{\frac{\eta_i}{2}}{\frac{\eta_i}{4}} x^{\frac{\eta_i}{4}}(\frac{1-\epsilon}{3})^{\frac{\eta_i}{4}}\right\}^{M}$

### S7. Base errors in DNA storage

The base-level error profiles including insertions, deletions, and substitutions are studied by analyzing data sets from four independent works (1, 4, 5, 12). And details of the used data sets are presented **in Supplementary S7 Table** 1. As shown in **Supplementary S7 Fig. 10**, the substitution error shows a much higher rate than insertion and deletion error rates regardless of the sequencing platforms. Using different sequencing techniques leads to varied error rates. Specifically, while taking less sequencing time, i.e., days versus tens days, Miseq-based systems turn to have ~3x higher base error rate than Hiseq-based systems. While both using the Hiseq-based system, the higher substitution error rate in the data set from (5) than the data set from (12) might due to the absence of the double PBSs and the higher ratio of non-overlap length against the total length, i.e., 0.42 vs 0.22.

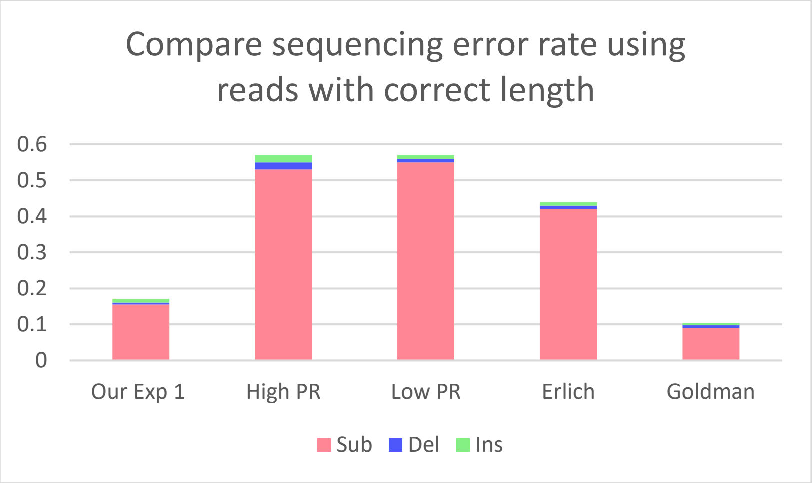


**Supplementary S7 Fig. 10 The comparison of sequencing error rate using reads with the correct length. Substitution errors dominate the error profile regardless of the used sequencing platforms.**

**Supplementary S7 Table 1 Details of the compared data set.**

| Data set | Sequencing company | Sequencing technique |
|---|---|---|
| High PR | Customa. | MiSeq |
| Low PR | Customa. | MiSeq |
| Erlich | Twist B. | MiSeq |
| Goldman | Agilent. | Hiseq |
| Our Exp 1 | Twist B. | Hiseq |

#### S8. 2/3/4-mer error patterns with errors occurring in the second and last positions

To comprehensively understand the patterns' effect on the error profile, in addition to the first position, errors that present in the second and last positions of the patterns are examined. The errors occurring in the second position is shown in **Supplementary S8 Fig. 11** while the errors occurring in the last position are shown in **Supplementary S8 Fig. 12**. From **Supplementary S8 Fig. 11**, it could be found that for each error type, the discrepancy among the patterns is much less significant than the discrepancy observed in patterns with the first position in errors as shown in Figure 3 in main text. In particular, for deletions in the 2-mer patterns (**Supplementary S8 Fig. 11A**), the homopolymer shows no distinguished higher error rate than other patterns. This might because that the alignment between the original sequence and the corrupted read (i.e., with a deletion in the homopolymer region) tends to align in a way that regards the base at the first position is deleted, i.e., allocating a gap mark at the first position. Hence, to effectively differentiate the homopolymer's effect on the deletion error rate from other patterns, the first position of the k-mer patterns should be taken as the observation point. For the 3-mer deletion patterns in **Supplementary S8 Fig. 11B**, all patterns with high error rates are tailed with the corresponding 2-mer homopolymer which is consistent with Figure 3A. The reason beneath is that the errors occurring at the second position of a 3-mer pattern could be also considered as errors occurring at the first position of a 2-mer pattern. Similarly, the 3-mer insertion patterns in **Supplementary S8 Fig. 11B**, all patterns with high error rates are tailed with the corresponding 2-mer patterns, i.e., AG, CG, TG, and GA which is consistent with the Figure 3A. Note that for 2-mer insertion patterns in **Supplementary S8 Fig. 11A**, the difference in error rates among patterns is quite insignificant. Compared with Figure 3A where the analyzed insertions occur in-between a 2-mer pattern, the observation of certain patterns with distinguished higher error further confirms that the neighboring bases in the original sequence have particular relevance to the probability of having an additional base inserted in the read (taken the original sequence as the reference). Again, same as substitution error patterns shown in Figure 3, there is no significant difference among patterns given that the second position as the observation anchor. As shown in **Supplementary S8 Fig. 12,** for all types of errors, the error patterns with the last position as the observation point show no prominent differentiation.

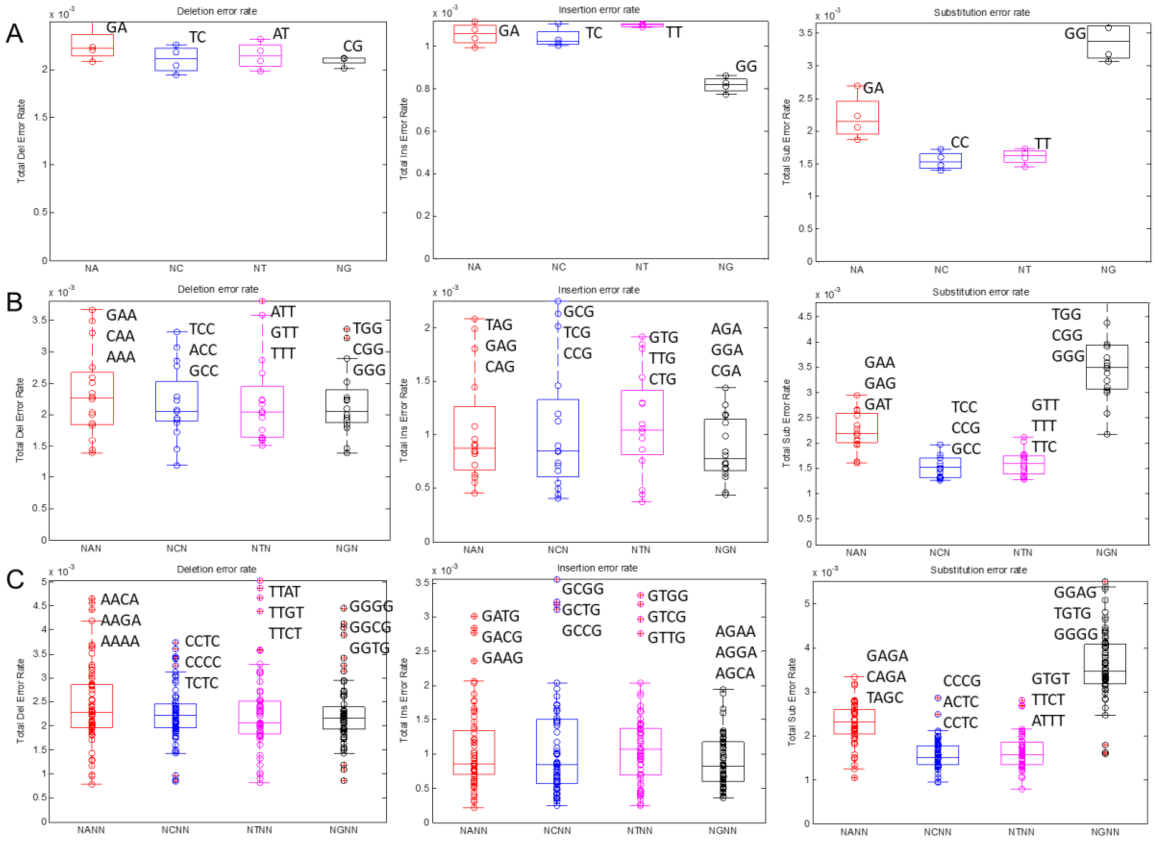


**Supplementary S8 Fig. 11 2/3/4-mer error patterns for deletions, insertions, and substitutions that occur in the second position of the pattern. The most erroneous patterns are marked correspondingly. (A) 2-mer patterns; (B) 3-mer patterns; (C) 4-mer patterns.**

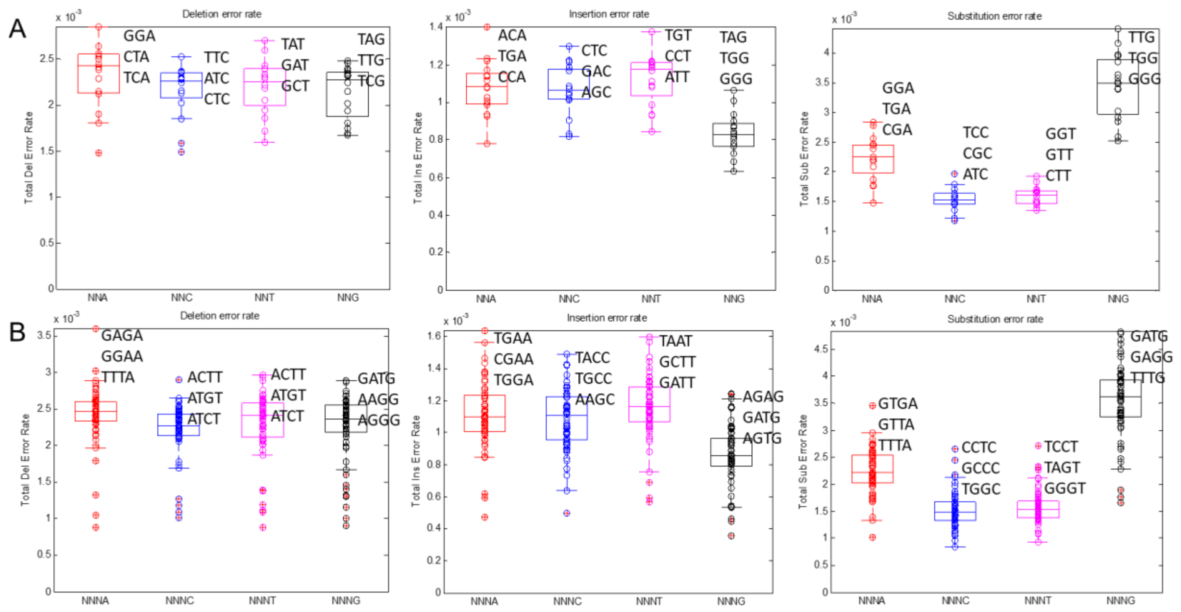


**Supplementary S8 Fig. 12 3/4-mer error patterns for deletions, insertions, and substitutions that occur in the last position of the pattern. The most erroneous patterns are marked correspondingly. (A) 3-mer patterns; (B) 4-mer patterns.**

#### S9. PCR stochasticity and PCR bias effects

The PCR amplification process could be seen as a stochastic branching process (10). In each cycle of PCR, the number of newly generated molecules follows a binomial distribution where the PCR efficiency is the underlying probability of the event that refers to the presence or absence of an offspring molecule copy. Considering that the PCR amplification is imperfect, i.e., PCR efficiency is less than 1.0, there exists stochasticity in the process. When the initial amounts of the molecule are small, the PCR stochasticity has a notable impact on the distribution of the copy number of the molecule. Following the iterative copy increment formulation $N(K+1) = N(K) + \mathbf{B}(N(K), p)$, where $\mathbf{B}(N(K), p)$ is a random variable of binomial distribution with parameters $N(K)$ and $p$; and $N(K)$ is the copy number at cycle $K$, we simulate the PCR process with small initial copies 1, 2, 3, and 4 under the conditions of PCR cycles 10 and PCR efficiency $p = 0.95$. As demonstrated in **Supplementary S9 Fig. 13**, in addition to one global maximum, there are other local maxima, which is therefore referred to as a multimode phenomenon. Moreover, the multimode phenomenon is more noticeable when the initial copy number is smaller.

Since sequences in single PBS data set have longer homopolymer, i.e., 4nt, which might introduce severer copy count bias compared with double PBS data set which only consists of sequences with homopolymer less than 3nt, the distribution bias difference between the two experiment data sets might be related to the PCR bias caused by the sequence-specific randomness. To further clarify whether PCR stochasticity or PCR bias is the main source of the distribution bias difference between the two count distributions, we used single PBS data set and separate it into sequences with 4nt homopolymer group (**Supplementary S9 Fig. 13B**) and sequences without 4nt homopolymer group (**Supplementary S9 Fig. 13C**). Compared with the distribution of all sequences in **Supplementary S9 Fig. 13A**, the spreads of distributions of two separated data sets have no significant difference, i.e., 2.75 versus 2.77 versus 2.69. Note that the percentage of references with 4nt homopolymer having 0 copy count is similar to the percentage of references with 4nt homopolymer in full sequence set, i.e., 76% versus 78%, implying there is no obvious increase of under-amplification for sequences with 4nt homopolymer. Moreover, the reference with maximum copy counts, i.e., 174, is with 4nt homopolymer, which further proves that the anticipation of lower amplification efficiency of sequence with longer homopolymer might not be the cause of distribution bias. These observations indicate that rather than distinct PCR bias caused by sequence-specific randomness (due to homopolymer differentiation), distinct degree of PCR stochasticity caused by varied sample preparation mainly explains the bias difference.

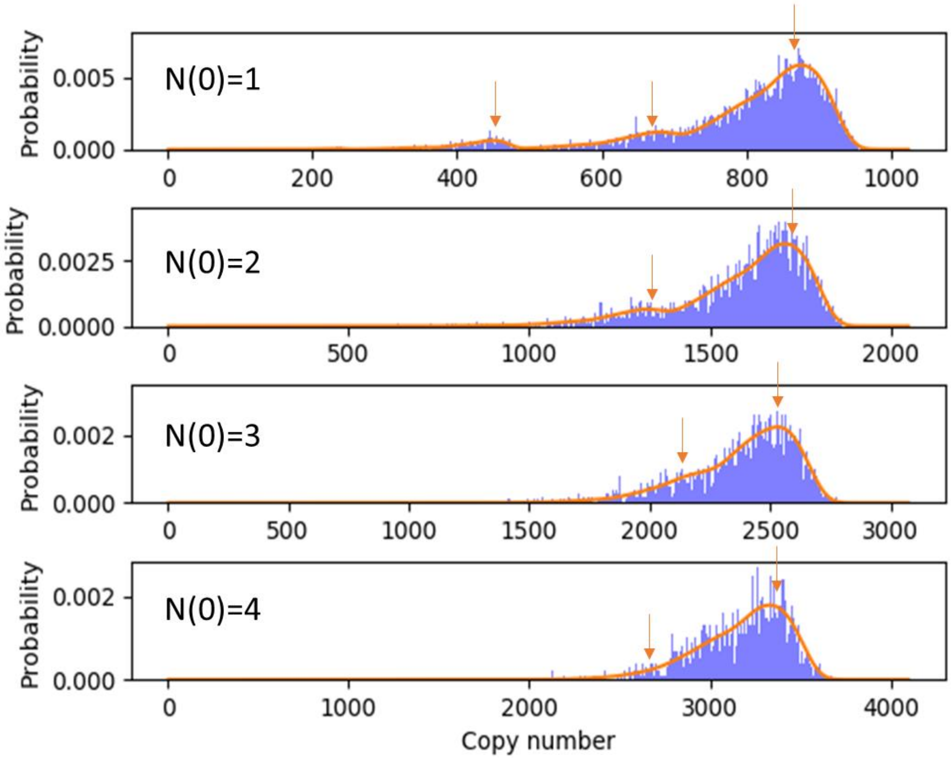


**Supplementary S9 Fig. 13 The probability density of the copy number distribution after 10 PCR cycles with PCR efficiency 0.95 and initial copy numbers of 1, 2, 3, and 4. Multimode phenomena in PCR amplification with low initial molecule copy amounts could be observed; and the lower the initial copy number, the more noticeable the multimode.**

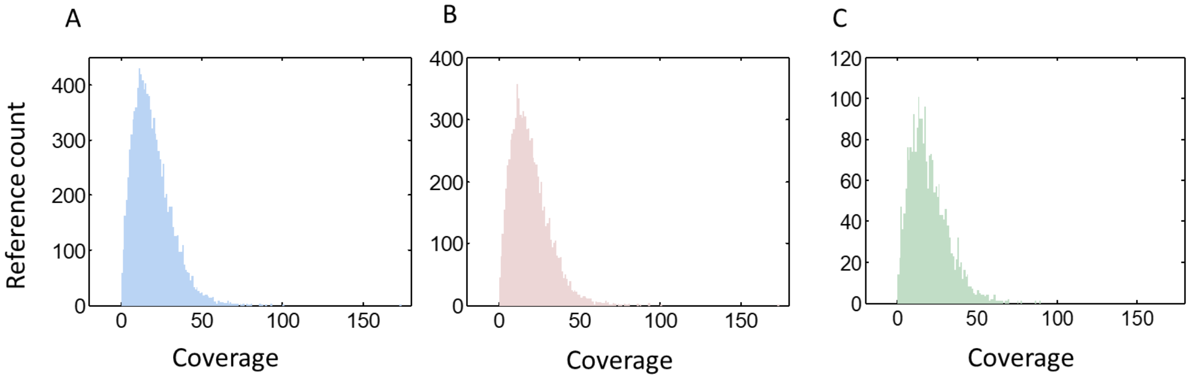


**Supplementary S9 Fig. 14 The comparison of copy count (or channel coverage) distributions of A. all sequences, B. sequences including 4nt homopolymer, and C. sequences without 4nt homopolymer with 20x coverage of single PBS data set.**

### S10. Data integrity relates to the structural design of the sequence

**Supplementary S10 Fig. 15** demonstrates the overall base error rates including substitutions, deletions, and insertions of four data sets from two different works (5, 6). The synthesis and sequencing techniques used in these two works are the same; but the sequence designs are varied. Specifically, the two compared systems correspondingly used fixed-length sequences appended with single PBS and variable-length sequences flanked with double PBSs as data unit. Comparing Exp 1(a) (from (5)) with Exp 2(a) (from (6)) in **Supplementary S10 Fig. 15**, it could be found that the overall base error rate is higher in the single PBS system than the double PBSs system. This might be since before sequencing there is only one additional molecule copy for each original synthesized single-stranded molecule that has been synthesized from one direction via the single strand complementation reaction in the single PBS system, leading to errors occurring in the latter stage of the base augmentation process could not be offset like double PBSs system where PCR amplification is applicable; and a large number of molecule copies are generated with augmentations proceed from two directions (taken the original molecule as reference). After filtering based on the read length, the overall base error rate of data set from (5), i.e., Exp 1(b), descends drastically, remaining substitutions as the major errors. However, for a variable-length system, i.e., Exp 2, reads could only be filtered in an inexact way, i.e., filtering upon a range of lengths, and therefore the insertions and deletions in the filtered data could not be reduced as much as Exp 1. In other words, while the high code efficiency and low complexity could be achieved, with the use of the variable-length design, the benefit of immediate filtering on resolving the indels at the receiver is restricted.

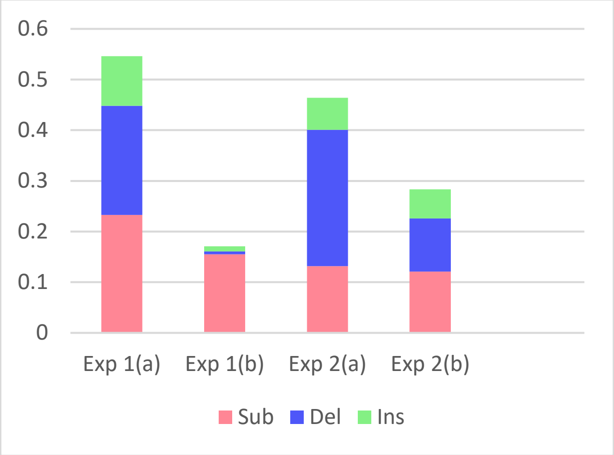


**Supplementary S10 Fig. 15 The comparison of base error rate between fixed-length single PBS and variable-length double PBSs DNA storage. The base error rates are analysed based on four data sets gained from two different works, i.e., Exp 1 and Exp 2, where (a) notation refers to data sets without filtering and (b) notation refers to data sets after filtering based on the length(s).**

### S11. The sequence error rate with or without filtering

As discussed above, filtering out reads with incorrect lengths could facilitate addressing insertions and deletions, alleviating the demand on the decoder. However, in some cases, filtering out these seemingly wrong reads might cause more sequence loss that requires more reconstruction efforts. To figure out whether filtering is beneficial to the decoder or not, we use the data from the fixed-length system (5) to conduct the analysis. From **Supplementary S11 Fig. 16A**, with a fixed number of reads, i.e., sequencing coverage, an increase in the sequence loss rate is observed when the reads with wrong lengths are filtered out. The trends of sequence loss rate decrease exponentially with the coverage, i.e., the read redundancy, which is consistent with the Poisson sampling effect that is widely observed in DNA sequencing. Meanwhile, as shown in **Supplementary S11 Fig. 16B**, the overall base error rate of the filtered reads is lower than the raw reads. Note that the base error rate is expected to be constant along the x-axis, i.e., sequencing coverage. However, the base error rates in the low coverage region, i.e., 1x~5x, are relatively larger than the plateau due to the bias of measure brought by insufficient samples from the population. In **Supplementary S11 Fig. 16C**, the overall sequence error rate that consists of sequence loss and sequence corruption (with the trial-and-error as the post-processing) are shown. It could be found that the overall sequence error rate of the filtered data set is higher than the rate of non-filtered data up to coverage 30x, suggesting that when the sequencing coverage, i.e., the number of redundant read copies is sufficient enough, while filtering might immediately reject erroneous data, the sequence loss caused by it should be carefully concerned especially for a highly biased data set, i.e., small size r or large dispersion 1/r.

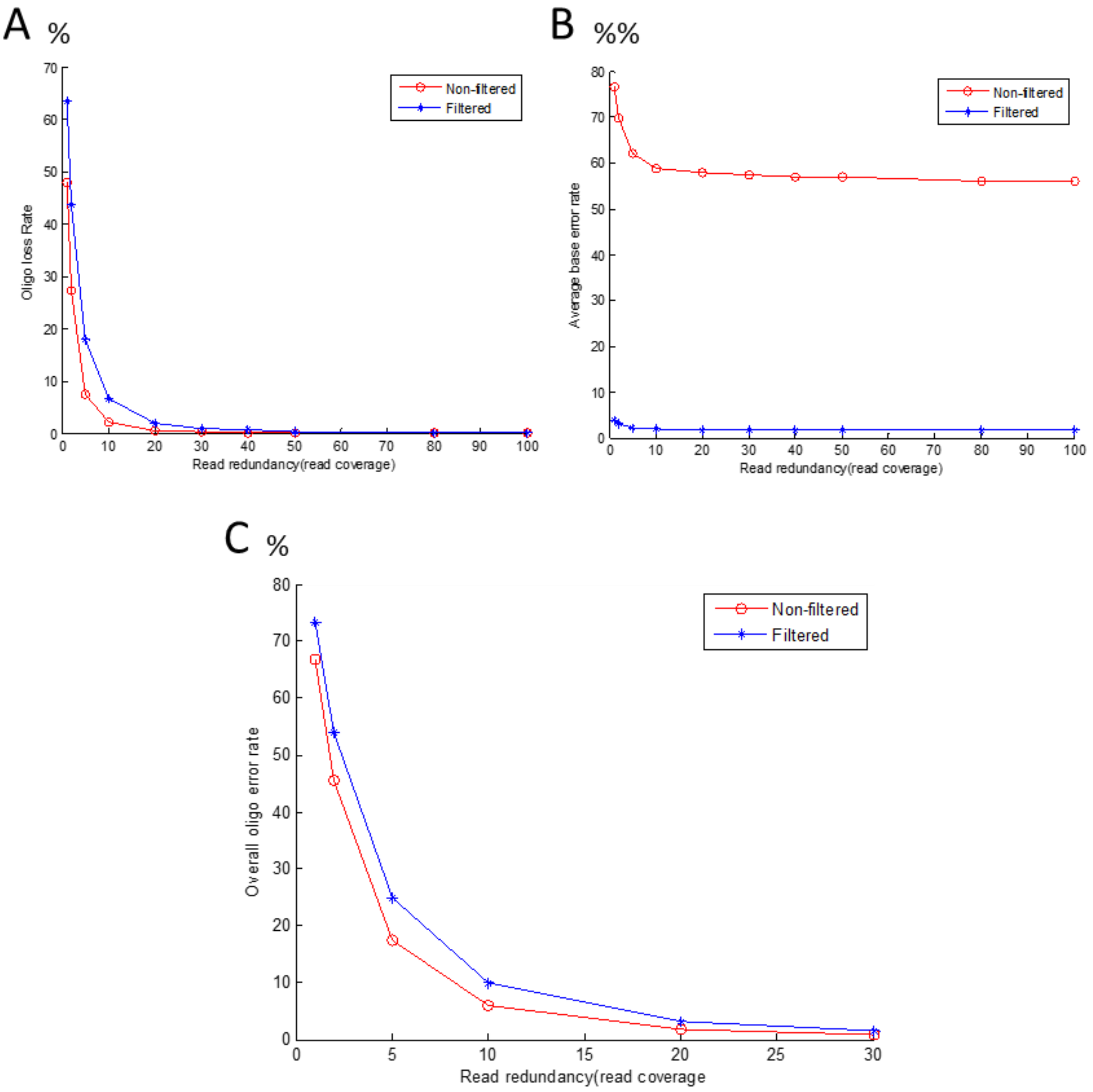


**Supplementary S11 Fig. 16 The filtering effect on the sequence loss and the overall sequence error rates. (A) The sequence loss rate against the sequencing coverage; (B) The base error rate; (C) The overall sequence error rate consisting of sequence loss and sequence corruption (trial-and-error) against the sequencing coverage.**

### S12. Theoretical analysis of the impact of sequence length

The relationship between the ratio of PE read length to the sequence length and the sequence corruption rate Ω follows,

$$\begin{cases} 1 & \frac{a}{b} < 0.5 \\ 1-(1-\varepsilon)^{2(b-a)}(1-\varepsilon/2)^{(2a-b)} & 0.5 \le \frac{a}{b} < 1 \\ 1-(1-\varepsilon/2)^{b} & \frac{a}{b} \ge 1 \end{cases}$$

where $a$ is the PE read length; $b$ is the sequence length; $\varepsilon$ is the raw base error rate. Specifically, when the ratio is below 0.5, stitching two PE reads is unable to recover the original sequence (as shown in **Supplementary S1 Fig. 1C**), leading to 100% corruption. When the ratio is from 0.5 to 1, the corruption rate decreases with the increase of the ratio. This is because the overlapped region is expanded when the ratio escalates. When the ratio is no less than 1, each PE read could ideally cover the whole sequence (**Supplementary S1 Fig. 1A**), rendering the merged read with a full overlapped region and consequently reducing the corruption rate to a low floor. The raw error rate used to draw Figure 6A in main text is set to 0.8% approximating the Illumina Miseq sequencing.

With PE read length $a$ fixed to 150, the sequence length $b$ is also observed to affect the sequence corruption rate and incremental redundancy for addressing the corruption. Particularly, the increased corruption rate caused by the increased sequence length requires an increased redundancy to recover data. The incremental redundancy follows $\frac{1}{1-\Omega}-1$, where $\Omega$ is calculated by $1-(1-\varepsilon)^{2(b-a)}(1-\varepsilon/2)^{(2a-b)}$, as $b \in (150, 300)$ which gives the ratio $\frac{a}{b} \in (0.5, 1)$.

Sequence length also has an impact on the achieved capacity. First, the redundancy required for correcting the sequence corruption lowers the achieved capacity. Second, the redundancy required for indexing is another paramount factor of the achieved capacity. These two redundancies are highly related to the sequence length; and impacts of sequence length on them are reverse. Specifically, given fixed PE read length, the longer the sequence length, the higher sequence corruption rate and error correction redundancy, and the lower achieved capacity of the system. In contrast, given fixed data size, the longer the sequence length, the lower number of sequences is required to store the data, the lower redundancy required to index the data, and thus the higher achieved capacity of the system. The achieved capacity with considerations of both error control correct redundancy and indexing redundancy could be derived as,

$$C = (1-\Omega) * (1 - b_i/b)$$

where $\Omega = (1-(1-\varepsilon)^{2(b-a)} * (1-\varepsilon/2)^{(2a-b)})^{\eta}$ is the sequence corruption rate counting in the effect of coverage $\eta$ with the assumption of the use of trial-and-error; other parameters are the same as the above mentioned; $b_i$ is the redundancy required to index all encoded sequences including information sequences and redundancy sequences. $b_i$ is resolved by equation,

$$4^{b_i} = \frac{M}{(b-b_i)(1-\Omega)}$$

where $M$ is the size of data that are to be stored.

**S13. Estimating sequence error rate when merged PE reads is used for decoding**

With the premise of using merged reads (from the PE reads) as the decoder's input, we theoretically estimate the reconstruction demand given different settings to the initial copy count distribution and base error rate. Specifically, the reconstruction demand on the decoder could be indicated by the sequence error rate at the receiver. The sequence error rate consists of sequence loss that relates to the initial copy count distribution and sequence corruption that relates to the base error errors. The initial copy count distribution depends on the synthesis techniques and the base error rate predominantly depends on the sequencing techniques. Moreover, given merging PE reads as the post-processing method after getting PE reads, the base error rate is no longer constant along the coordinate of the sequence, instead, it varies subject to the characteristic of the region it belongs to, i.e., overlapped and non-overlapped regions. Considering that the overall base error rate of the merged reads is affected by the sequence length which decides the ratios of the overlapped and non-overlapped regions, two extreme cases, i.e., the merged reads with fully overlapped (main text Figure 7) and fully non-overlapped regions (**Supplementary S13 Fig. 17**), are separately considered. Build upon these considerations and with the assumption of using trial and error before decoding, we formulate the estimation of overall sequence error rate as follows,

$$(\frac{r}{\eta+r})^r + \left(1 - (\frac{r}{\eta+r})^r\right) * (1 - (1-\varepsilon)^{2(b-a)} * (1-\varepsilon/2)^{(2a-b)})^\eta,$$

where $r$ is the size parameter of the copy count distribution (which approximates to negative binomial (NB) distribution at the receiver; $\eta$ is the mean coverage of reference sequence at the receiver; $\varepsilon$ is the raw base error rate (i.e., error rate in non-overlapped region); $b$ is sequence length with range $a$ to $2a$ where $a$ is the PE read length. Note that the formulation separately considers sequence loss which is deduced by size parameter $r$ and mean coverage $\eta$ and sequence corruption which is deduced by raw base error rate $\varepsilon$, sequence length $b$, PE read length $a$, and mean coverage $\eta$. Another more precise way to construct estimation formulation is using random variable coverage $\eta_i$ rather than mean coverage $\eta$ where $\eta_i$ follows NB distribution $NB(r, p = \frac{r}{\eta+r})$. In this way, the sequence loss ($\eta_i = 0$) and sequence corruption ($\eta_i \neq 0$) can be denoted simultaneously. To demonstrate the estimation in different scenarios, values of parameters used in the formula are empirically set according to practice. For example, in main text Figure 7 and **Supplementary S13 Fig. 17**, size parameter $r$ is set from 1 to 7 according to the synthesis technique and biological process; raw error rate $\varepsilon$ is set to 0.8% and 0.3% according to the Illumina Miseq and Hiseq sequencing platforms; PE read length $a$ is set to 150 according to the commonly used PE150 sequencing protocol; mean coverage $\eta$ is set from 1 to 30 for illustration; while sequence length $b$ is set differently, for main text Figure 7, $b$ is set to 300 for one extreme case where the merged reads with fully overlapped region and , $b$ is set to 150 in **Supplementary S13 Fig. 17** for the other extreme case where the merged reads with fully non-overlapped region.

**Supplementary S13 Fig. 17** demonstrates estimation results of the fully overlapped case where the sequence length equals the PE read length. Overall, the higher the base error rate is, the higher the sequencing redundancy/coverage is required to achieve the equivalent sequence error rates. While with two different base errors in the two sub-figures, the representations of the curves are quite similar owing to the low sequence corruption rate leveraged by the short sequence length and the fact that the merged reads are fully overlapped. Besides, it could be universally found that the smaller size parameter r is, i.e., the more over-dispersion of the copy count distribution, the higher the sequence error rate (due to the higher sequence loss rate), suggesting that concerns on the impact of copy count distribution bias on reconstruction demand are indispensable.

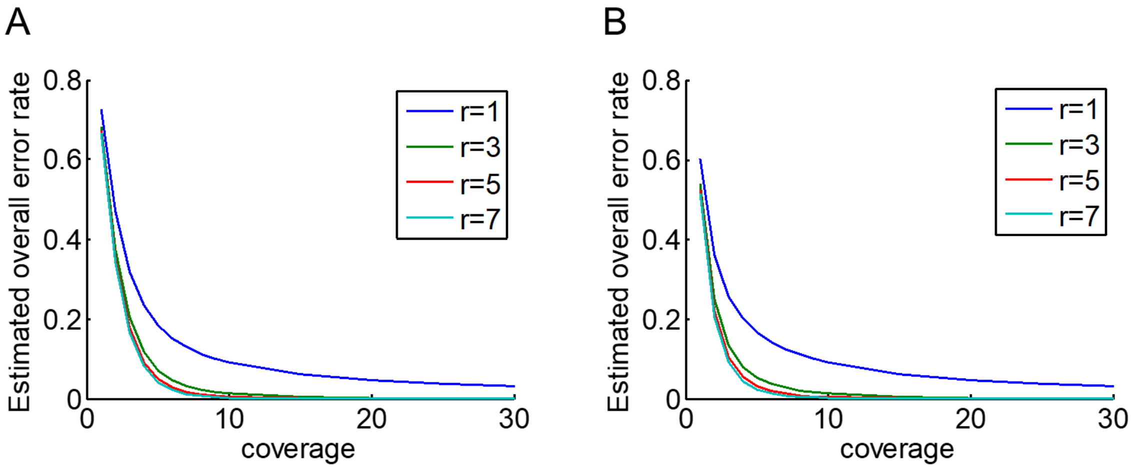


**Supplementary S13 Fig. 17 Theoretical estimation of the overall oligo error when the merged reads from the PE reads are fully overlapped. In this case, the sequence length equals the read length, i.e., 150nt. According to different sequencing techniques, the raw base error rates are set to (A) 0.8% similar to Miseq; (B) 0.3 similar to Hiseq. Curves with different colours represent different initial copy count distributions.**
